# Exposure to Mites Sensitizes Intestinal Stem Cell Maintenance, Splenic Marginal Zone B Cell Homeostasis, And Heart Development to Notch Dosage and Cooperativity

**DOI:** 10.1101/2020.02.13.948224

**Authors:** Francis M. Kobia, Kristina Preusse, Quanhui Dai, Nicholas Weaver, Praneet Chaturvedi, Sarah J. Stein, Warren S. Pear, Zhenyu Yuan, Rhett A. Kovall, Yi Kuang, Natanel Eafergen, David Sprinzak, Brian Gebelein, Eric Brunskill, Raphael Kopan

## Abstract

Cooperative DNA binding is a key feature of transcriptional regulation. Here we examined the role of cooperativity in Notch signaling by CRISPR-mediated engineering of mice in which neither Notch1 nor Notch2 can homo- or heterodimerize, essential for cooperative binding to sequence paired sites (SPS) located near many Notch-regulated genes. While most known Notch-dependent phenotypes were unaffected in Notch1/2 dimer-deficient mice, a subset of tissues proved highly sensitive to loss of cooperativity. These phenotypes include heart development, compromising viability in combination with low gene dose, and the gut, developing ulcerative colitis in response to 1% DSS. The most striking phenotypes – gender imbalance and splenic marginal zone B cell lymphoma – emerged in combination with dose reduction or when challenged by chronic fur mite infestation. This study highlights the role of the environment in malignancy and colitis, and is consistent with Notch-dependent anti-parasite immune responses being compromised in the dimer deficient animals.

**Highlights:**

- Notch dimerization has an *in vivo* role in contributing to intestinal homeostasis
- Loss of cooperativity can manifest as Notch gain or loss of function phenotypes
- Mite infestation exacerbates all phenotypes, triggers MZB hyperproliferation in mutant animals
- Mite-infested mutant mice develop SMZL with age

## Introduction

The evolutionarily conserved Notch receptors and ligands influence metazoan development and adult tissue homeostasis by directly translating an *inter-*cellular interaction into *intra-*cellular transcriptional outputs that control cell fate, proliferation, differentiation, and apoptosis (Artavanis-Tsakonas et al., 1999; Bray, 2006; Kopan and Ilagan, 2009). Mammals possess four Notch receptors (N1 to N4) and five Delta/Jagged ligands; all of which are Type I transmembrane proteins. The Notch pathway stands out relative to other signaling pathways in lacking signal amplification: canonical Notch signaling is initiated when a ligand on one cell engages a Notch receptor on a neighboring cell. This interaction unfolds the receptor’s juxtamembrane region enabling cleavage by the metalloprotease ADAM10. The truncated, cell membrane bound polypeptide is then cleaved by the γ-secretase complex freeing the Notch Intracellular Domain (NICD), which subsequently translocates into the nucleus (Kopan and Ilagan, 2009; Kovall et al., 2017). NICD associates with the DNA-binding protein CSL (<u>C</u>BF1/<u>S</u>uppressor of Hairless/<u>L</u>AG-1, also known as RBPJ in vertebrates) and recruits the coactivator Mastermind; thereby assembling a <u>N</u>otch <u>T</u>ranscription <u>C</u>omplex (NTC) that activates Notch target gene expression (Bray, 2016; Gordon et al., 2008; Kovall et al., 2017).

The Notch pathway plays complex and context-dependent roles during development and adult tissue homeostasis. Perturbations in the Notch pathway are associated with developmental syndromes (Masek and Andersson, 2017) and cancers (Aster et al., 2017; Siebel and Lendahl, 2017). For example, *Notch1* promotes T-cell development (Pui et al., 1999; Radtke et al., 2010; Radtke et al., 1999), while *Notch2* is indispensable for <u>M</u>arginal <u>Z</u>one <u>B</u>-cell (MZB) development (Gibb et al., 2010; Hozumi et al., 2004; Liu et al., 2015; Maillard et al., 2004; Moran et al., 2007; Saito et al., 2003; Tanigaki et al., 2002; Witt et al., 2003). Accordingly, elevated Notch1 signaling is oncogenic in T-cells driving Acute Lymphoblastic Leukemia (T-ALL) (Ellisen et al., 1991; Weng et al., 2004) while increased Notch2 signaling is associated with splenic marginal zone B-cell transformation (Arcaini et al., 2016; Piris et al., 2017; Santos et al., 2017). Inversely, where Notch signals promote differentiation, the pathway can have a tumor suppressor function with diminished Notch signaling being associated with cancer (Demehri et al., 2009; Koch and Radtke, 2007).

How can the Notch pathway control multiple, dissimilar outcomes? Because each Notch receptor is consumed as it generates a signal and cannot be reused, signal strength has emerged as a key in controlling outcome. Indeed, some Notch-dependent processes are exquisitely sensitive to dosage and manifest both haploinsufficient and triplomutant effects in tissues such as the fly wing (Bray, 2006; Guruharsha et al., 2012). In mammalian tissues, signal strength, defined as the sum of NICD released from all ligand-bound Notch receptors on the cell surface, is a far more important determinant of Notch signaling outcome than NICD composition (i.e. N1ICD vs N2ICD) (Liu et al., 2015; Liu et al., 2013; Ong et al., 2006). Several mechanisms have been found to modulate signal strength including receptor glycosylation (Haltiwanger, 2008; Kakuda and Haltiwanger, 2017), force-generating ligand endocytosis (Gordon et al., 2015; Langridge and Struhl, 2017), the contact interface between cells (Shaya et al., 2017), and the ratio of receptor/ligand constitution within the cell (Sprinzak et al., 2010). More recently, ligand-dependent signal dynamics were demonstrated to be another key determinant of signaling outcome (Nandagopal et al., 2018).

Interestingly, NICD can assemble into dimeric, cooperative NTCs (Arnett et al., 2010) on <u>S</u>equence <u>P</u>aired <u>S</u>ites (SPSs), first described in the regulatory regions of the *Drosophila* E(spl) (enhancer of split) locus (Bailey and Posakony, 1995). SPSs consist of two DNA binding sites orientated in a head to head manner (Bailey and Posakony, 1995; Nellesen et al., 1999) separated by 15-17 nucleotides (Nam et al., 2007). NICD dimerization is facilitated via a conserved interface in NICD’s ANK (Ankyrin repeat) domain. In human N1ICD, this interface consists of Arg^1985^, Lys^1946^ and Glu^1950^. Dimerization in Notch1 is effectively abolished by mutating Arg^1985^ into Ala^1985^ (R^1985^A), Lys^1946^ into Glu^1946^ (K^1946^E) or Glu^1950^ into Lys^1950^ (E^1950^K), and loss of dimerization results in reduced activation of dimer-dependent targets (Arnett et al., 2010; Liu et al., 2010a; Nam et al., 2007).

Given the conservation of the dimer interface in most Notch receptors (*C. elegans* being the exception), the conservation of SPSs near known Notch targets (Hass et al., 2015; Liu and Posakony, 2012, 2014), and the ability of synthetic SPSs to regulate NICD levels (Kuang et al., 2019), we hypothesized that the precise mixture of agnostic and dimer-sensitive targets in a given cell will couple with Notch signal strength and dynamics to shape the responses to Notch signals and contribute to the context-specificity of Notch-related pathologies. To test this hypothesis in a mammalian species, we evaluated mice homozygous or hemizygous for dimerization-deficient alleles of *Notch1* (*N1*^*RA*^) and *Notch2* (*N2*^*RA*^) and found that dimerization contributes to context-specific Notch activity. We report that in the hemizygous state, dimer deficient Notch molecules are haploinsufficient in the heart and intestine, with lethal consequences when modified by the presence of the ectoparasite, *Demodex musculi* (fur mites). Mice homozygous for a dimer deficient *N1*^*RA/RA*^ allele displayed a female-biased, mild cardiac phenotype consistent with Notch loss-of-function. Conversely, *N2*^*RA/RA*^ mice displayed a striking over proliferation of marginal zone B cells in parasite-infested mice, but not in mite-free mice. In time, the cell-type specific gain of function activity of *N2*^*RA/RA*^ produced a splenic marginal zone lymphoma-like phenotype in mite-exposed mice. Mechanistically, these effects reflect a shift in target amplitude favoring monomer-driven targets, and reduced negative feedback.

## Results

### Generation of N1 and N2 dimerization deficient mice

To interrogate the physiological role of dimerization, we first quantified the effective binding cooperativity of NTC to SPS and CSL sites using purified RBPj, the N1 RAM-ANK domain, and a MAML protein in electrophoretic mobility shift assays (EMSA; see supplemental information). We compared the mobility of NTC complexes containing either wild type N1ICD or a mutant version with the substitution of a single critical Arginine residue (Arg^1974^) located in the mouse N1 ANK domain to alanine (Figure 1A-B, Supplemental Figure S1A-D). Quantifying the intensity levels of the bands in the image and fitting the data to a binding model that takes into account cooperative binding (see supplemental methods), we measured the cooperativity contributed by NTC binding to either an SPS probe (1xSPS) or a CSL probe containing two sites (2xCSL). We find that both wild type and mutant NTC had no cooperative binding to the 2xCSL probe. When the 1^st^ site was already bound, the wild type NTC displayed a 5-fold stronger binding to the 2^nd^ site in the 1xSPS probe compared to the binding to the 1^st^ site. Importantly, NTC containing N1^RA^ICD had no cooperative binding to the 2^nd^ site in the 1xSPS probe (Fig. 1A, Supplemental Figure S1A-D).

**Figure 1.**
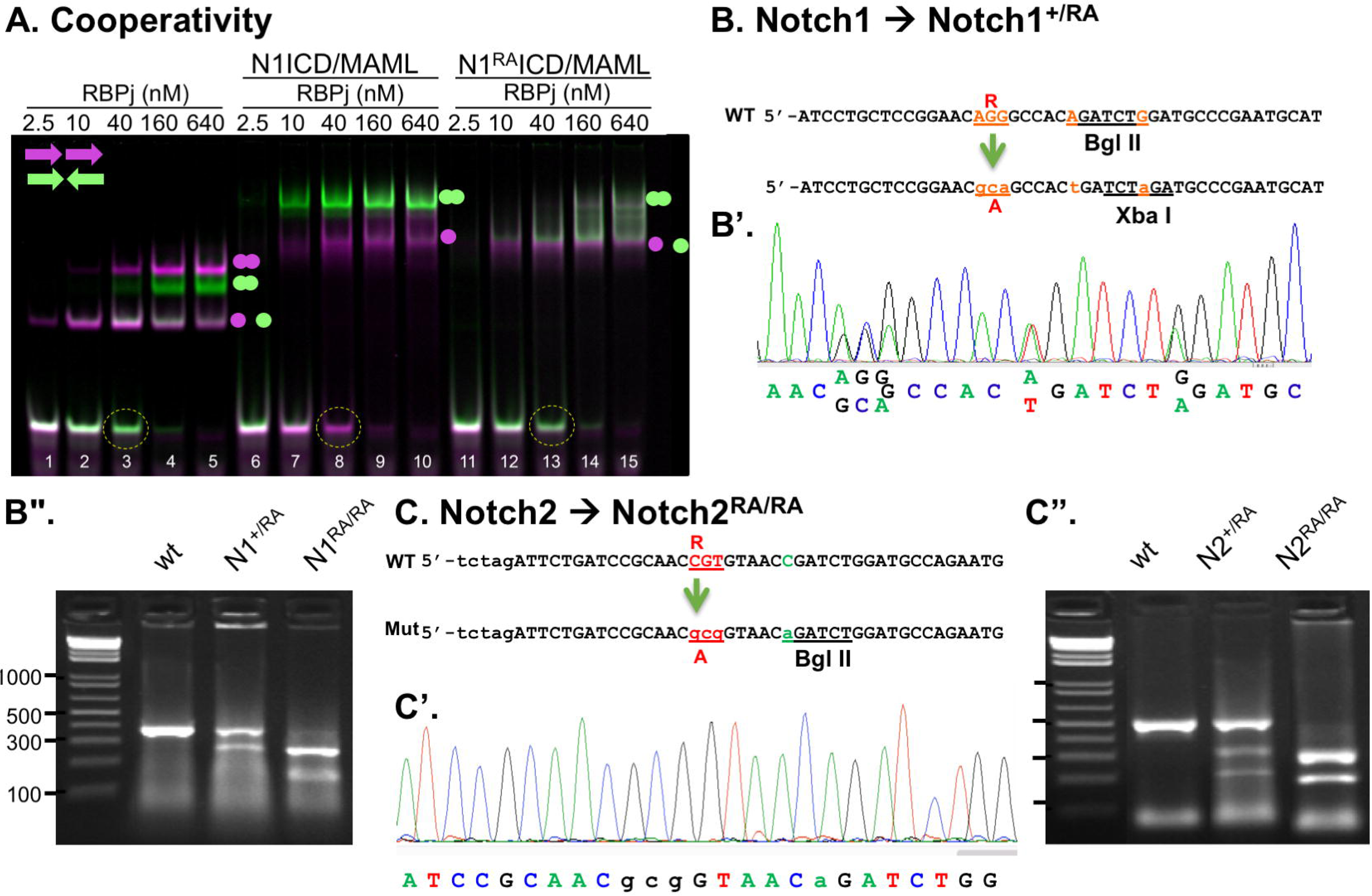
Generation of Notch dimerization deficient mice. (A) Electrophoresis mobility shift assay for purified proteins binding to CSL (magenta) or SPS (green) probes. Balls mark occupancy of one or two sites. Note that cooperative binding by wild type NTC, but not the RA mutant NTC, specifically depletes the SPS probe, but not the CSL probe (compare lanes 8 and 13). See text and Supplemental Figure S1 for detail. (B-C) CRISPR-Cas9-mediated double-strand break was used to mediate homologous recombination of a short oligonucleotide into Exon 32 substituting Arg *N1*^*R1974*^ (B-B”) and *N2*^*R1934*^ into Ala (C-C”), generating *N1*^*RA/RA*^ and *N2*^*RA/RA*^ animals. To facilitate genotyping, two silent mutations (in red) were included in the oligo, abolishing the BglII restriction site while generating an XbaI site in *Notch1* (B’). In Notch2, a silent mutation (in green) was included to create a BglII site (C’). Sequencing PCR products containing these regions confirmed the presence of the Arg to Ala substitution in founders; digestion of these PCR products with XbaI (for N1, B”) and BglII (for N2, C”) confirmed the presence of one (N1) or two (N2) mutant alleles.

To ask if loss of cooperativity impacts function *in vivo*, we used CRISPR-Cas9 to introduce amino acid substitutions at Arg^1974^ and Arg^1934^ in N1 and N2, respectively. Following the generation of a double strand break in Exon 32 of N1 or N2, a short oligonucleotide harboring two linked mutations was homologously recombined. Nucleotides coding for Arg^1974^ in N1 (N1R^1974^) and Arg^1934^ in N2 (N2R^1934^) were changed to code for Ala, creating the N1^R1974A^ (N1^RA^) and N2^R1934A^ (N2^RA^) mutations, respectively (Figure 1B, E). To facilitate genotyping of N^RA^ animals silent mutations were introduced to abolish a BglII restriction site while creating an XbaI restriction site in N1^RA^ (Figure 1B), or to generate a BglII restriction site in N2^RA^ (Figure 1E). Animals that carry one (N1^+/RA^ or N2^+/RA^) or two (N1^RA/RA^ or N2^RA/RA^) mutated chromosomes were first identified by digesting PCR products of the targeted chromosomal regions in Exon 32 with XbaI and/or BglII. XbaI cuts the N1^RA^ chromosome but not the wild type PCR product; BglII cuts the *N2*^*RA*^ but not the N2 wild type PCR products (Fig1 D, G). The presence of each respective RA mutation in *N1*^*RA*^ and *N2*^*RA*^ founders was subsequently confirmed by direct DNA sequencing (Fig1 C, F). Animals deficient for N1 dimerization (*N1*^*RA*/*RA*^), N2 dimerization (*N2*^*RA/RA*^) or both N1 and N2 dimerization (*N1*^*RA/RA*^; *N2*^*RA/RA*^) were generated by crossing founders. All were viable, fertile and without any overt phenotype at P360 in mixed or C57BL6 background (unless otherwise stated, all subsequent results were generated in the mixed background). Because Notch1 signaling plays a central role in T-cell development (Radtke et al. 1999, 2010), we expected that loss of Notch1 dimerization would negatively impact the T-cell compartment. However, an analysis of the T-cell compartments in the thymus and spleen as well as thymic T-cell sub-compartments (DN, SP and DP), revealed a normal T-cell compartment in N1^RA/RA^ mice relative to wild type controls (Supplemental Figure S2). This finding and the normal lifespan of *N1*^*RA/RA*^; *N2*^*RA/RA*^ animals, was surprising since we anticipated many dimerization-dependent genes (e.g., *Nrarp, Hes1, Hes5* and *Myc*; (Arnett et al., 2010; Hass et al., 2015) would be negatively impacted as seen when constitutively active wild type and dimer mutant Notch proteins lacking the extracellular domain (NΔE and N^RA^ΔE) are overexpressed in cell culture (Supplemental Figure S1E).

### Homologous *N1*^*RA/RA*^; *N2*^*RA/RA*^ mice display barrier defects in the colon and reduced proliferation in the crypt when challenged with 1% DSS

Notch1 and Notch2 act redundantly during the development and maintenance of the gut to block Klf4-induced niche exit of intestinal stem cells (Pellegrinet et al., 2011; Riccio et al., 2008), and Math1-induced differentiation of secretory cells (Noah and Shroyer, 2013; Shroyer et al., 2007). However, no pathology was revealed in histological examination of *N1*^*RA/RA*^; *N2*^*RA/RA*^ intestines (see below). Several studies have shown that a chronic decrease in Notch signaling compromised the intestinal barrier and exacerbated colitis in different mouse models (Mathern et al., 2014; Obata et al., 2012; Okamoto et al., 2009). Under the assumption that a compromised barrier may increase sensitivity to dextran sulfate sodium (DSS)-induced colitis, we asked if intestinal homeostasis and intestinal barrier were robust enough in *N1*^*RA/RA*^; *N2*^*RA/RA*^ by exposing mice to DSS. When given 2.5% DSS in their drinking water, wild type mice developed colitis as evident by a mild weight loss whereas co-housed *N1*^*RA/RA*^; *N2*^*RA/RA*^ mice exhibited severe weight loss necessitating euthanizing within one week (Figure 2A). To ask if *N1*^*RA/RA*^; *N2*^*RA/RA*^ were predisposed to develop colitis we exposed them to 1% DSS, which does not affect the weight of control littermates even after repeated exposures (Figure 2B, C). By contrast, co-housed *N1*^*RA/RA*^; *N2*^*RA/RA*^ mice experience significant weight loss, some had blood in stool at end of the third and fourth DSS cycle, and one had to be removed during the first cycle of DSS following severe weight loss and blood in stool. However, all remaining mice recovered well when DSS was removed from the drinking water, even after multiple cycles of treatment (Figure 2B). Histological analysis of the colons revealed injury in wild type mice and a complete disruption of colonic epithelium of *N1*^*RA/RA*^; *N2*^*RA/RA*^ mice treated with 2.5% DSS (Figure 2D, top). After four periods of 1% DSS exposure *N1*^*RA/RA*^; *N2*^*RA/RA*^ mice had a more severe injury than controls or 2.5% DSS exposed wild type mice (Figure 2D). Relative to baseline (Figure 2E), proliferation in wild type crypt epithelia (Krt8/18 positive) was elevated in 1% or 2.5% DSS treated controls (Figure 2F). By contrast, we noticed a surprising decrease in proliferation competence (less Ki67) (and mitosis (Phospho-H3 positive cells)) in colonic epithelium following the fourth 1% DSS treatment in crypts of *N1*^*RA/RA*^; *N2*^*RA/RA*^ mice (Figure 2G) relative to wild type or untreated crypts, suggesting a role for Notch cooperativity within colonic stem cells recovering from injury. Finally, RA-specific while changes in barrier markers were not detected, a significant increase in IL4, Th17A and IL1ß was observed in DSS-treated *N1*^*RA/RA*^; *N2*^*RA/RA*^ relative to wild type (Figure 2H) consistent with enhanced inflammatory response driving colitis in these animals. Thus, these results are consistent with *N1*^*RA/RA*^; *N2*^*RA/RA*^ alleles being hypomorphic loss-of-function alleles in maintaining intestinal homeostasis and immune response.

**Figure 2.**
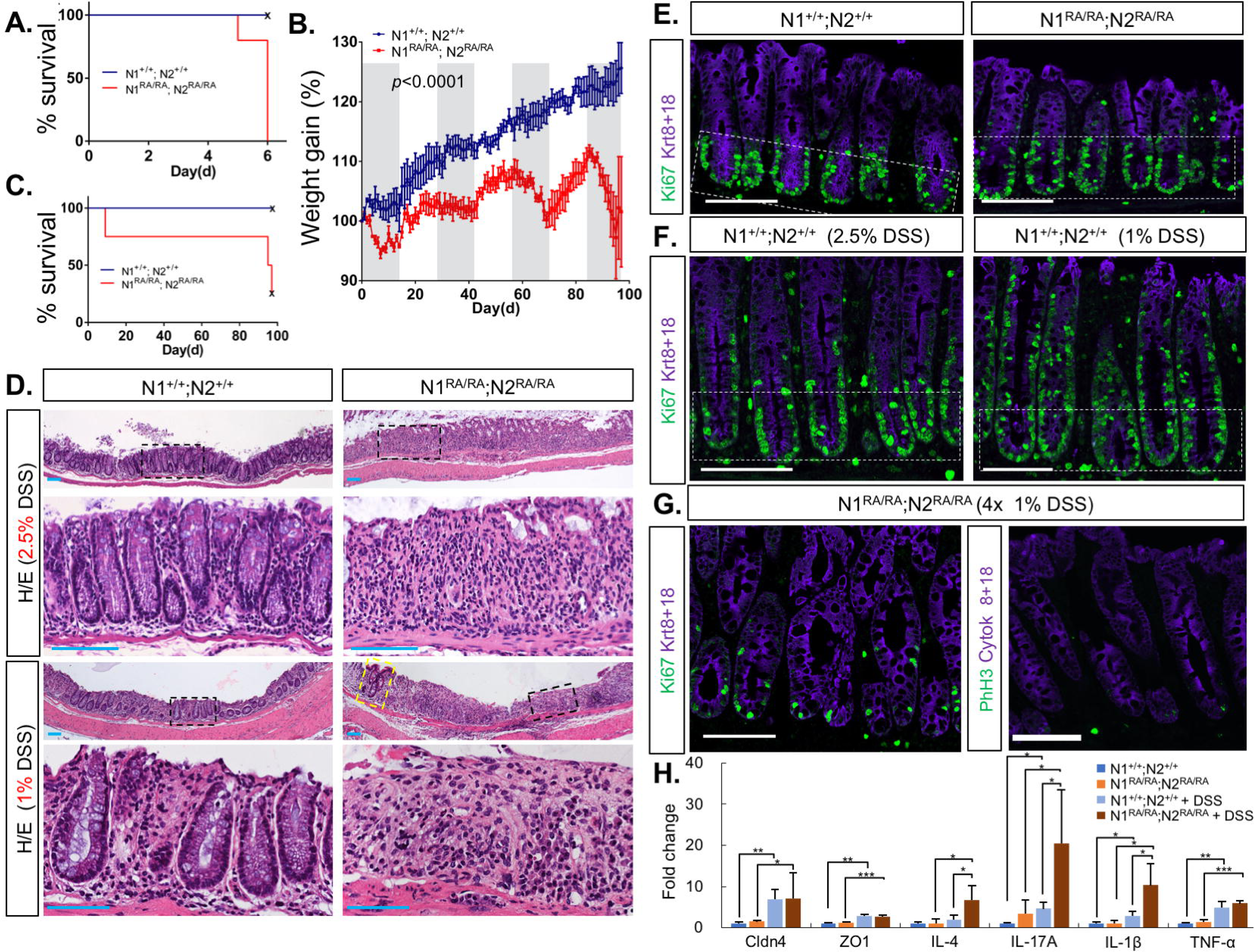
Notch dimerization deficient mice are sensitized to DSS-induced colitis. (A) All *N1*^*RA/RA*^; *N2*^*RA/RA*^ mice exposed to 2.5% DSS treatment had to be euthanized due to severe weight loss before day seven. (B) Daily weight measurements of wild type (blue) or mutant (red) mice treated with alternate cycles of 1%DSS (gray sections) or no DSS (white sections). (C) Survival curve of 1% DSS treated mice. Three of four mutant mice had to be euthanized due to sever weight loss by the fourth cycle. (D) Hematoxylin-eosin staining of colonic tissue from DSS treated mice. Dashed black boxes are enlarged below. Yellow dashed box showed a section with crypts in an otherwise injured colon in 1% DSS treated *N1*^*RA/RA*^; *N2*^*RA/RA*^ mice. (E) Ki67 staining of untreated mice with the indicated genotypes, crypt regions (Ki67+) are boxed. (F) Increased proliferation competence in colon of DSS treated wild type mice (Krt8/18+, Ki67+ staining outside the box). (G) Decreased proliferation competence in colon of *N1*^*RA/RA*^; *N2*^*RA/RA*^ mice exposed to four rounds of 1% DSS. (H) qPCR on RNA extracted from distal colon of *N1*^*RA/RA*^; *N2*^*RA/RA*^ and WT mice treated for 11 days with 1% DSS; n=3. (*, *p*<0.05).

### N1^RA^; N2^RA^ hemizygotes display impaired intestinal cell proliferation and fate allocation in the crypt and cause ventricular septum defects in the heart

The wild type and R-A mutant monomeric NICD protein bind CSL sites similarly (Figure 1A, Supplemental Figure S1), supporting the hypothesis that the signal generated in *N1*^*RA/RA*^; *N2*^*RA/RA*^ mice activated sufficient non-SPS gene targets to a level needed to support Notch-dependent decisions. If the Notch RA alleles are hypomorphic, challenging R-A mutant mice by lowering protein levels might reveal additional phenotypes. To test this idea, we generated an allelic series in which one copy of *N1* or *N2* was deleted in the *N1*^*RA/RA*^; *N2*^+*/*+^ or *N1*^+*/*+^; *N2*^*RA/RA*^ backgrounds, respectively, as well as deleting one copy of *N1* and one copy of *N2* deleted in the *N1*^*RA/RA*^; *N2*^*RA/RA*^ background. Single N1 or N2 RA hemizygotes resembled R-A mutants; however, crossing *N1*^*RA/RA*^; *N2*^*RA/RA*^ with *N1*^+*/–*^; *N2*^+*/–*^ animals generated significantly fewer *N1*^*RA/–*^; *N2*^*RA/–*^ pups at birth than expected (*p*<3×10^−9^, Chi-squared distribution analysis, Supp Table S1).

*N1*^+*/–*^; *N2*^+*/–*^ double hemizygous animals have a normal lifespan and no overt phenotype under normal housing conditions, but have been reported to develop mild cardiac phenotypes (Garg et al., 2005; Nigam and Srivastava, 2009). Given the critical role the circulatory system plays during gestation, we suspected that the poor representation of *N1*^*RA/–*^; *N2*^*RA/–*^ pups at birth was due to defects in cardiac development. Analysis of E16.5 embryos born to mite-free dams revealed highly penetrant and severe Ventricular Septal Defects (VSD) in the hearts of *N1*^*RA/–*^; *N2*^*RA/-*^ embryos (Figure 3), compromising viability. Milder VSD was also observed with lower penetrance in N1^RA/-^; N2^+/RA^ embryos (Figure 3). Around this time, the colony became infected with fur mites, but the infestation was not immediately detected on the sentinels. Strikingly, when we retroactively segregated the fecundity data in our colony management software, we observed a significant (*p=*0.02, χ^2^) skewing of the gender ratio in C57BL6 *N1*^*RA/RA*^ carriers, with male:female ratio nearing 2:1 in fur mite infested mice, as opposed to the normal 1:1 ratio in mite-free mice (Supp Table S2).

**Figure 3.**
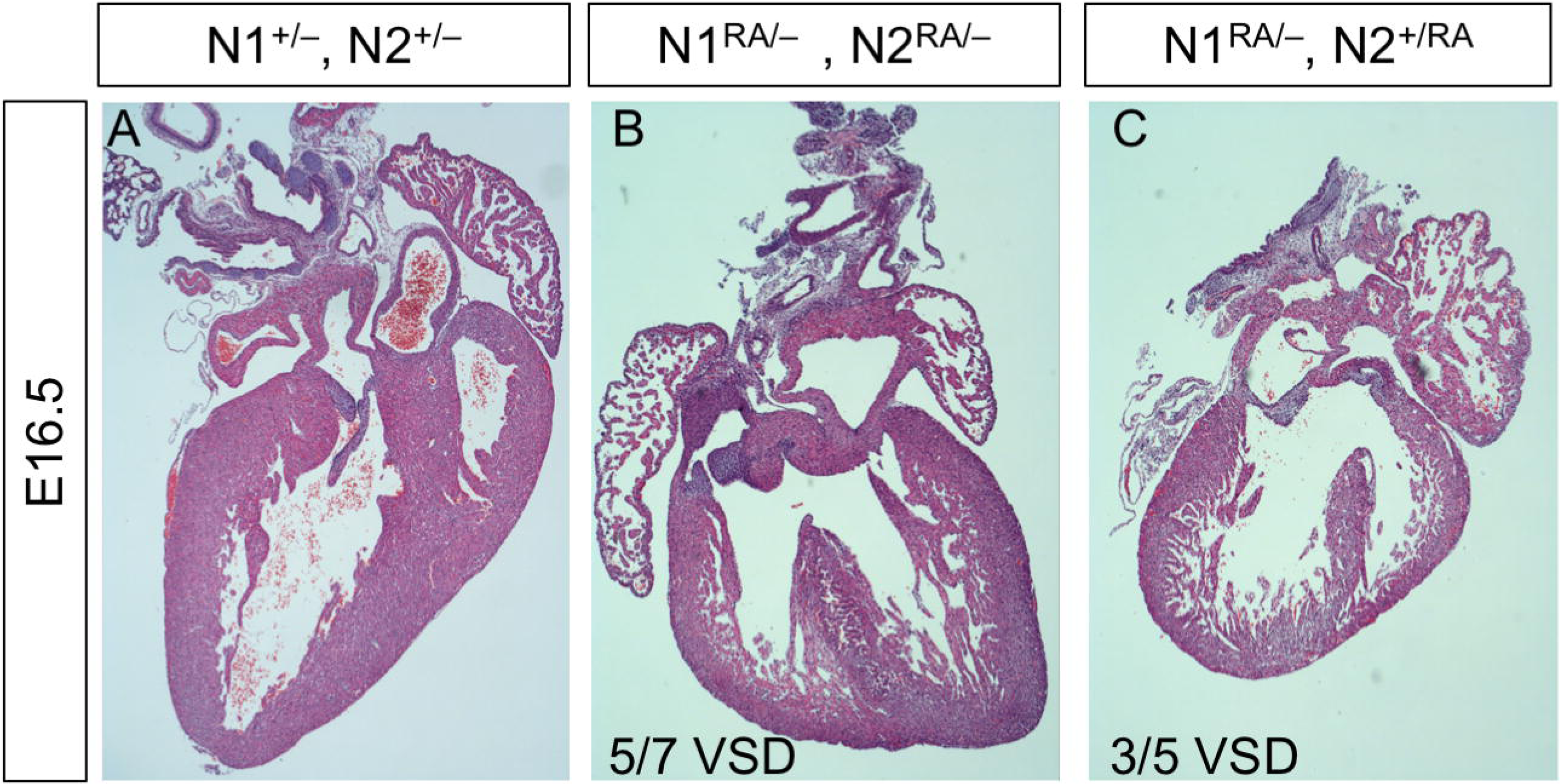
Notch1^R1974A^ substitution compromised ventricular septum development. (A) Normal heart with complete septum. (B) *N1*^*RA/–*^; *N2*^*RA/–*^ heart with a severe, highly penetrant VSD. The penetrance by age shown below the image. (C) *N1*^*RA/–*^; *N2*^+*/RA*^ heart with a milder, less penetrant VSD.

The longest-lived *N1*^*RA/–*^; *N2*^*RA/–*^ pup born to mite infested, mixed background dams was much smaller than its litter mates and was sacrificed at P30 when it became moribund. Necropsy revealed a striking and complete loss of the intestinal crypts and villi (Carulli et al., 2015; Riccio et al., 2008; van Es et al., 2005; Vooijs et al., 2007). Since complete failure to form an intestine would have not permitted survival outside the womb, we assumed this phenotype reflected loss of intestinal stem cells (Figure 4A, A’) consistent with the loss of Ki67 in DSS-challenged *N1*^*RA/RA*^; *N2*^*RA/RA*^ mice (Figure 2G). To investigate this phenotype further, we assessed proliferation and fate allocation in the developing intestine of E18-P1 animals with various allele combinations. To examine fate allocation, we stained sections with alcian blue and analyzed the number of secretory Goblet cells, which are known to expand in Notch hypomorphic background (Noah and Shroyer, 2013; Vooijs et al., 2007). The small intestines of control or *N1*^+*/–*^; *N2*^+*/–*^ P0 pups were indistinguishable (Figure 4B, C). By contrast, alcian blue staining revealed a significant expansion in the goblet cell compartment at P0 in *N1*^*RA/–*^; *N2*^*RA/–-*^ pups born to fur mites infested dams (Figure 4D). In mice free of mites, the overall morphology appeared normal with few regions displaying excess goblets. To assess proliferation, we stained adjacent intestinal sections with antibodies against Ki67 and Phospho-H3. Strikingly, whereas trans-hemizygote *N1*^+*/–*^; *N2*^+*/–*^ intestines resembled wild type (Figure 4E-F), the *N1*^*RA/-*^; *N2*^*RA/-*^ intestine from mite-infested dames contained very few Ki67 positive cells (Figure 4G-H). We also noted a reduction in the number of cells undergoing DNA replication in particular, and competence to self-renew in general, in *N1*^+*/–*^; *N2*^*RA/RA*^ intestines (Supplemental Figure S3C-C’, G,G’). However, upon eradication of *Demodex musculi* with permethrin, all E18.5 and all but 1/8 surviving P0 *N1*^+*/–*^; *N2*^+*/–*^ pups had at least some regions within the small intestine in which Ki67-positive tissue could be observed (Figure 4J, the percentage of Ki67+ crypts; Supplemental Figure S3I-J’; note reduced proliferation in Ki67-positive crypts of a P1 pup post-mite eradication). Overall, these data reflect an impaired ability to maintain intestinal stem cells in *N1*^*RA/–*^; *N2*^*RA/–*^ mice, which was exacerbated by *Demodex musculi* infestation. Because these phenotypes can reflect complex interaction between multiple tissues in the *N1*^*RA/RA*^; *N2*^*RA/RA*^ and *N1*^*RA/–*^; *N2*^*RA/–-*^ mice, we deferred the mechanistic analysis of this gene-environment interaction in ISC maintenance to future investigation.

**Figure 4.**
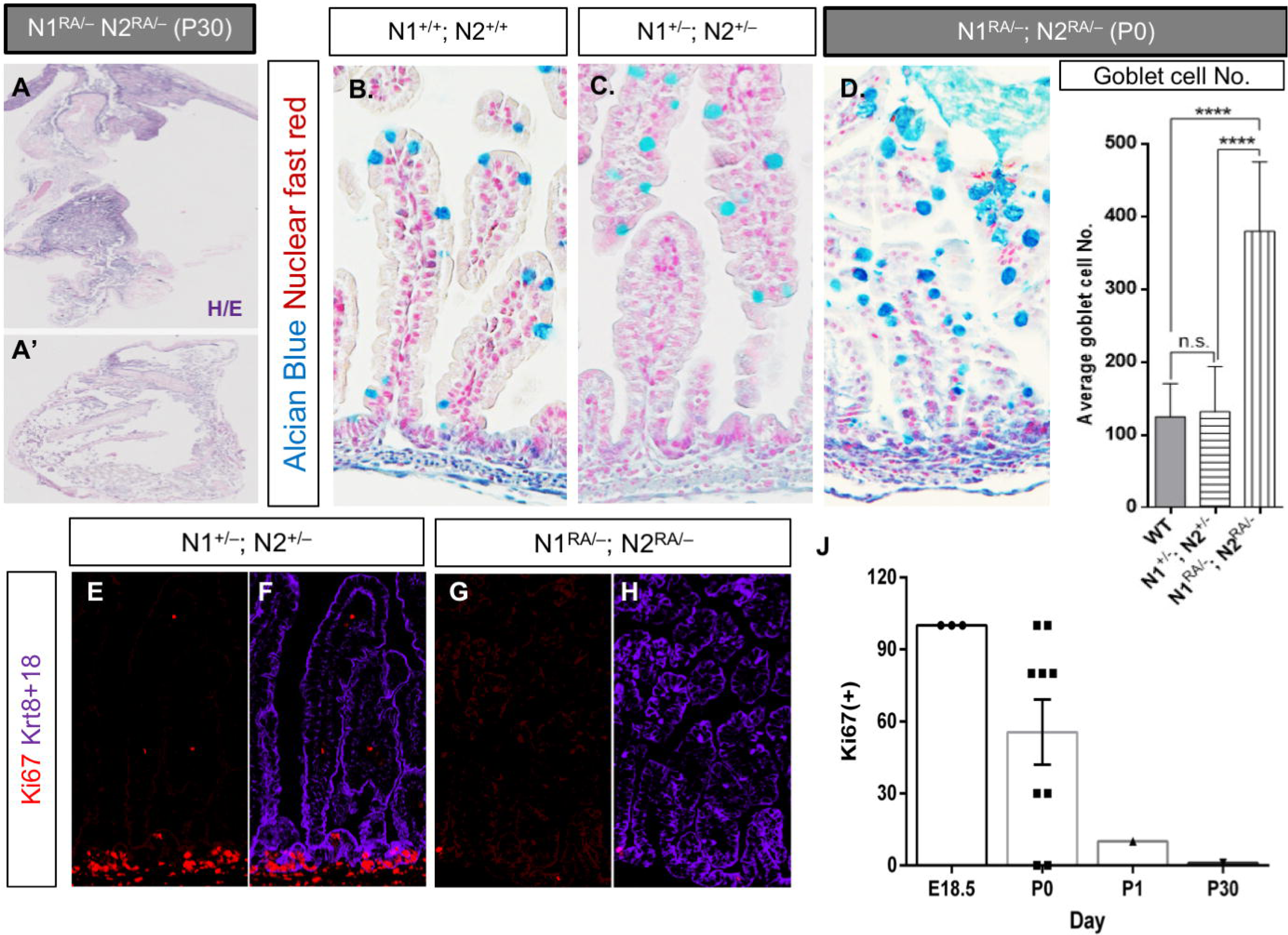
Notch dimerization deficient mice are hypomorphic for Notch activity in the gut. (A-C) Alcian blue staining of *N1*^*RA/RA*^; *N2*^*RA/RA*^ and *N1*^+*/–*^; *N2*^+*/–*^ adult intestines. (D) Deletion of one N1 and one N2 allele in the RA background (*N1*^*RA/–*^; *N2*^*RA/–*^) caused lethality; loss of the intestine (presumably due to stem cell exhaustion) was evident in the longest surviving pup at P30. (E-G) Alcian blue analysis of P0 intestines detected an increase in goblet cell numbers in *N1*^*RA/–*^; *N2*^*RA/–*^ (G) relative to wild type or *N1*^+*/–*^; *N2*^+*/–*^ intestines (E-F), quantified in (H). (I-J) Ki67 staining in P0 intestine from control *N1*^+*/–*^; *N2*^+*/–*^ (I) or *N1*^*RA/–*^; *N2*^*RA/–*^ (J) newborn. (K) quantification of Ki67-positive area at the indicated age. Among surviving *N1*^*RA/–*^; *N2*^*RA/–*^ pups during (n=2) and after (n=12) fur mite infestation.

### The N2^RA^ allele drives expansion of MZ B-cells in *Demodex musculi* infested mice

Marginal zone B-cells (MZB) are exquisitely sensitive to *N2* dosage (Gibb et al., 2010; Hozumi et al., 2004; Liu et al., 2015; Maillard et al., 2004; Saito et al., 2003; Tanigaki et al., 2002; Witt et al., 2003). Even a mild reduction in N2ICD can generate a noticeable decline in MZB cell numbers (Liu et al., 2015). We used FACS to analyze the splenic B cell population and asked if *N2*^*RA*^ displayed haploinsufficient phenotypes in the dose-sensitive MZ compartment. Intriguingly, we observed a significant increase in the MZB compartment in *N2*^*RA/RA*^ spleens in a mixed-background mouse exposed to *Demodex musculi* (Figure 5A). The size of other splenic B-cell subsets, including the immediate MZB precursors (MZP), Follicular B-cells (FoB) and transitional type-2 cells (T_2_) did not change significantly in either *N2*^*RA/RA*^ (Figure 5A), *N2*^+*/–*^ or *N2*^*RA/–*^ (Figure 5B), inconsistent with a fate switch.

**Figure 5.**
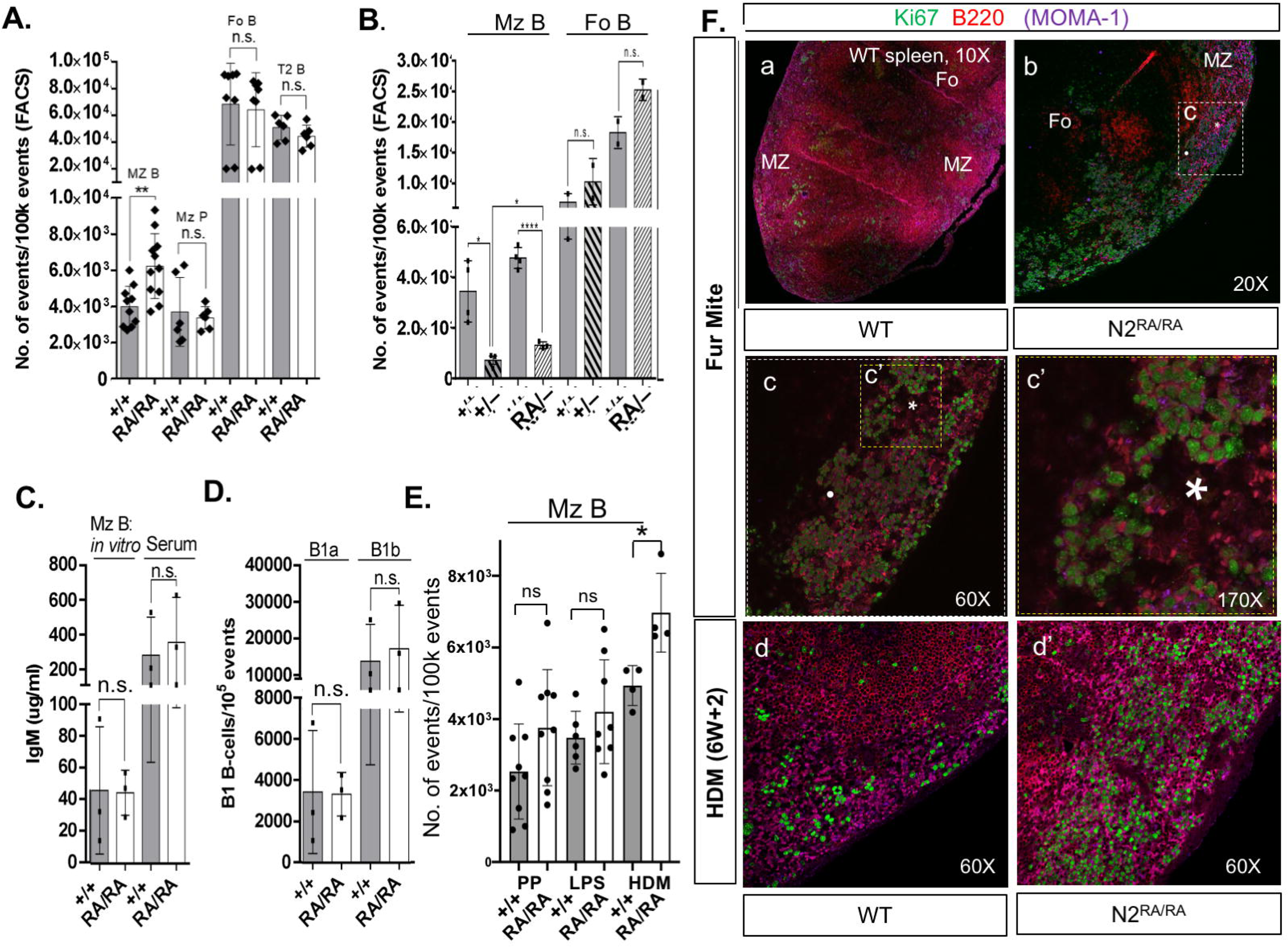
Loss of Notch2 dimerization expands the splenic MZ B-cell compartment in the presence of mites. (A) FACS analysis of *N2*^*RA/RA*^ fur mite infested mouse spleens revealed an expansion the MZ B-cell compartment relative to wild type (+/+). Values for MZ P, F0, and T2 splenic B-cell populations were not significantly different; n=11. (B) The MZB-cell compartment in heterozygotes. (C) LPS-induced differentiation and IgM production *in vitro* and serum levels of IgM *in vivo;* n=4. (D). Peritoneal B1a B-cell compartment in mite infested wild type and *N2*^*RA/RA*^ mice. (ns=not significant, n=2). (E). The MZB-cell compartment in *N2*^*RA/RA*^ mice after treatment with permethrin, an immunosuppressant used to manage fur mite infestation (PP), after LPS, or after 6 weeks of treatment with house dust mites (HDM) extract. (*, *p*<0.05; ns=not significant). In (A-E) P-values generated by two-tailed t-test. Error bars represent s.d. (F). Proliferation in the splenic marginal zone of mite infested mice: (Fa) The spleen from a mite infested control animal or (Fb) from *N2*^*RA/RA*^ mice. Boxed region magnified in Fc and again, in Fc’, asterisk and dot provided for orientation. Magnification noted. (Fd, d’) 60x view of marginal zone from control and HDM treated *N2*^*RA/RA*^ mice.

MZB cells reside in the splenic marginal zone where they surveil for blood-borne pathogens. Upon encountering antigens, they rapidly differentiate into IgM producing plasmablasts that secrete vast amounts of IgM before undergoing apoptosis (Cerutti et al., 2013). To test IgM levels *in vivo*, we performed ELISA on serum collected from unstimulated *N2*^*RA/RA*^ and littermate controls and detected no significant difference in the levels of circulating IgM (Figure 5C). To test for the possibility that loss of N2 dimerization negatively impacted differentiation into IgM producing plasmablasts upon stimulation, we FACS isolated MZB and FoB cells from mite-exposed *N2*^*RA/RA*^ and wild type spleens. We then stimulated these cells *in vitro* with LPS for 5 days followed by imaging to assess for proliferation, and by ELISA to quantify the IgM levels secreted into the medium. Relative to wild type MZB cells, there was no appreciable difference in the ability of N2^RA/RA^ MZB cells to respond to LPS stimulation (Supplemental Figure S4A), and ELISA analysis detected no significant difference in the amount of IgM secreted by *N2*^*RA/RA*^ MZB cells relative to wild type controls (Figure 5C). Finally, we tested if loss of dimerization affected the development of B1 B cells, which predominantly reside in the peritoneal cavity (Witt et al., 2003). FACS analysis revealed no significant changes in the levels of either the B1a or B1b subsets of B1 B-cells between *N2*^*RA/RA*^ and wild type animals (Figure 5D). Together, these data suggest that MZB cell numbers did not increase due to a fate switch or a failure to differentiate properly, and are consistent with a proliferative phenotype triggered by the N^RA^ mutation.

In the course of these experiments, the *Demodex musculi* infestation was eradicated with permethrin. Within a few months, as we saw with other phenotypes, mite-free *N2*^*RA/RA*^ animals no longer showed MZB expansion (Figure 5E, post-permethrin, PP). Since permethrin has been reported to suppress the immune system in mice (Blaylock et al., 1995; Punareewattana et al., 2001), we assumed permethrin transiently suppressed MZB proliferation. Multiple subsequent analyses over many months, however, failed to detect a difference in MZB numbers between *N2*^*RA/RA*^ and control mice post eradication and permethrin treatment, suggesting that this allele was indistinguishable from wild type in untreated and pathogen free animals. Accordingly, we compared MZB numbers between *N2*^+*/–*^ spleens, which have decreased MZB numbers due to haploinsufficiency, with *N2*^*RA/–*^ spleens and found no further reduction in MZB numbers (Liu et al., 2015; Saito et al., 2003). These data suggest that the *N2*^*RA*^ has neither a haploinsufficient character for the *N2*^*RA*^ allele in the MZB linage (Figure 5B), nor a proliferative phenotype triggered cell-autonomously by the N2^RA^ protein. To ask if environmental stimuli interacted with N2^RA^ within the spleen, mice were exposed to bacterial lipopolysaccharide (LPS) or to dermatitis, produced by twice weekly exposure of the ear epidermis to dust mite extract (HDM, *Dermatophagoides farinae*) for 6 weeks (see methods). MZB cell numbers were analyzed 8 weeks after the first LPS injection or HDM extract application. LPS induced transient IgM production but did not change MZB census in the spleen at 8 weeks (Figure 5E, LPS). Importantly, HDM-induced dermatitis elevated MZB numbers only in the *N2*^*RA/RA*^ mice but not in controls (Figure 5E, HDM). Collectively, these experiments establish that specific gene-environment interactions produce the MZB phenotype in *N2*^*RA/RA*^ mice.

To assess the rate of splenic B-cell proliferation in mite-infested mice, we stained C57BL/6J *N2*^*RA/RA*^ and WT spleen sections with antibodies against Ki67, B220 (to mark the B-Cells) and Moma-1 (to mark the marginal zone). We detected defuse B220 staining with elevated Ki67 in marginal zone cells in the *N2*^*RA/RA*^ spleen (Figure 5F), as well as in germinal centers (rich in FoB-cells and strongly B220 positive, Supplemental Figure S4), not seen in WT spleen of co-housed animals (Supplemental Figure S4B-D). Caspase 3 activation was not observed in *N2*^*RA/RA*^ sections (not shown). Increased marginal zone proliferation in *N2*^*RA/RA*^ relative to wild type control was also detected in spleens after six weeks of HDM treatment (Figure 5Fd, d’). Combined, these data suggest that mite infestation or prolonged HDM exposure leads to enhanced MZB proliferation.

### Aged *N2*^*RA/RA*^ mice have enlarged spleens but only *Demodex musculi* infested mice progress to a Splenic Marginal Zone B-cell Lymphoma (SMZL)-like state

As noted above, we observed no change in lifespan in *N2*^*RA/RA*^ or *N1*^*RA/RA*^; *N2*^*RA/RA*^ animals. However, as old animals were culled, we noticed enlarged spleens in animals carrying the *N2*^*RA/RA*^ allele post mite eradication (Supplemental Figure S5A). Spleens from mice infested by *Demodex musculi* were much larger, the most severe splenomegaly exhibited by *N2*^*RA/RA*^; *N1*^+*/RA*^ (Figure 6A-E), perhaps because they contained a higher contribution of the inflammation-prone FVB strain (Quigley et al., 2011; Quigley et al., 2009). Histologically, spleens from mite-infested *N2*^*RA/RA*^ carriers lost their typical architectures with *N2*^*RA/RA*^; *N1*^+*/RA*^ spleens showing the greatest disruption. Splenic morphology showed an expansion of the white pulp at the expense of the red pulp. In the most severe cases, the spleens appeared to consist of white pulp only (Figure 6F-K). In addition, two *N2*^*RA/RA*^; *N1*^+*/RA*^ animals had multiple enlarged abdominal lymph nodes. H/E staining revealed a striking morphological similarity to the enlarged spleens from the same animals, suggesting infiltration by a splenic population, or lymphoproliferation within the LN (Figure 6L-L”). To further characterize the nodes, we stained the enlarged spleens and lymph nodes for the B-cell markers B220 and Pax5. Both were positive for the B-cell markers (Figure 6M-N’). These features are reminiscent of human SMZL (Splenic Marginal Zone B-cell - Lymphoma), a clonal B-cell neoplasm that manifests itself in elderly patients and consists of small lymphocytes that infiltrate the lymph nodes and other organs (Arcaini et al., 2016; Piris et al., 2017; Santos et al., 2017). As would be expected, the enlarged lymph nodes and spleens were highly proliferative as indicated by Ki67 and phospho-H3 staining (Supplemental Figure S5B,D). Paradoxically, human SMZL are associated with hyperactive NOTCH2 signaling due to an N2ICD-stabilizing truncation upstream to the WSSSP sequence in the PEST domain of NOTCH2 (Kiel et al., 2012; Peveling-Oberhag et al., 2015; Rossi et al., 2012), whereas the *N2*^*RA*^ alleles are hypomorphic in expression assays and in the context of the heart, crypt, and gut barrier. To confirm that this line of CRISPR-modified mice did not inherit a truncated *Notch2* allele, we sequenced Exon 34 DNA isolated form lymph node filled with B-cells and found a perfect match to the wild type *Notch2* allele (ENSMUSG00000027878).

**Figure 6.**
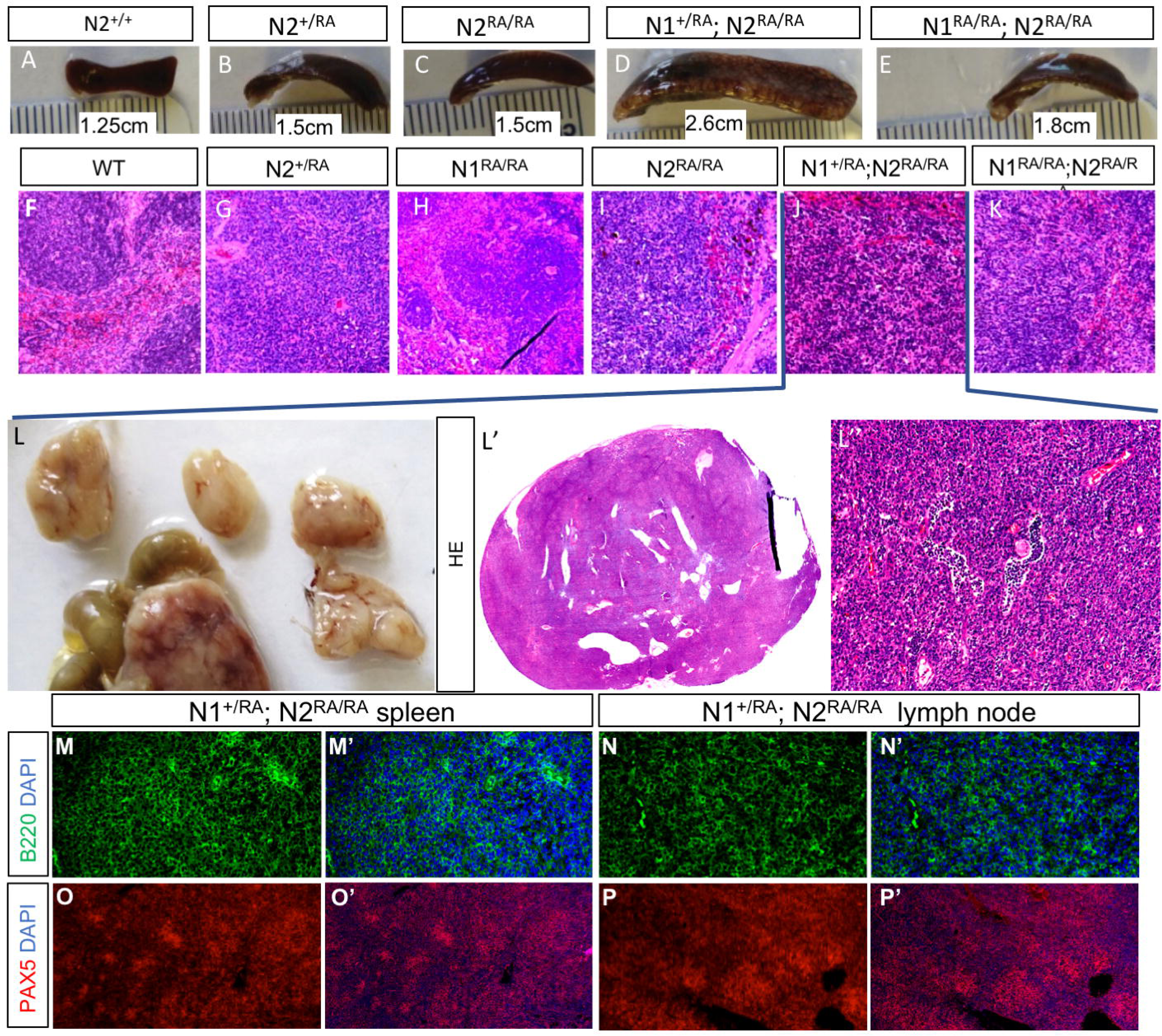
Aged *N2*^*RA/RA*^ and *N2*^*RA/RA; N1*+*/RA*^ mice develop severe splenomegaly and tumors reminiscent of Splenic Marginal Zone B-cell Lymphoma (SMZL). (A-E) Images of spleens from mice P600-P700 with indicated genotypes. (F-K) Severely enlarged spleens are associated with disorganized morphology and the appearance of cells with large nuclear:cytoplasmic ratio (genotypes indicated, see low magnification image in Supplemental Figure S6E). (L) large enteric lymph nodes detected in the individual shown in (J), densely populated with cells with large nuclear:cytoplasmic ratio (L’-L”). Staining the spleen (M) and lymph node (N) with Ki67 and the B-cell markers B220 and Pax5 identifies infiltrating cells as spleen-derived B cells. Magnification: 10x.

### Loss of Notch dimerization in *Demodex musculi* infested mice did not stabilize N2ICD in MZB cells but promoted their proliferation of MZB cells via the Myb-FoxM1 axis

Recently, we identified an unanticipated consequence of increasing the number of SPS sites in the *Drosophila* genome, namely, accelerated degradation of phospho-NICD (Kuang et al., 2019). Further, we demonstrated that this enhanced degradation affects some, but not all, Notch dependent decisions in *Drosophila*. Bristle precursor cells requiring a pulse of Notch were refractory to the destabilizing effect, whereas wing margin cells reliant on prolonged Notch signals were sensitive to NICD (Kuang et al., 2019). To test if the similarity of our phenotype to SMZL might in part be reflective of N2ICD stabilization, we assessed N2ICD levels in nuclear extracts from wild type and *N2*^*RA/RA*^ MZB cells. We did not detect a significant accumulation of nuclear N2ICD in sorted *N2*^*RA/RA*^ MZBs (Supplemental Figure S6), suggesting that the gain-of-function phenotypes observed in *N2*^*RA/RA*^ MZB cells is not due to enhanced N2ICD stabilization. Thus, unlike in many human SMZL patients, increased N2ICD stability is unlikely to be the cause of the SMZL phenotype in the N2^RA/RA^ mice.

In *N2*^*RA/RA*^ mice, Notch pathway activity in T-cells and skin is intact as it relies on *Notch1*. Assuming the MZB effect is cell autonomous we addressed the mechanism involved in enabling a haploinsufficient allele to drive MZB expansion via transcriptome and chromatin analyses. We sorted and performed RNAseq on MZB cells from spleens of three *N2*^*RA/RA*^ and three wild type littermates infested with *Demodex musculi*. The data were mapped to the mouse genome (mm10), and PCA analysis indicated that one of the triplicates in each group was an outlier. To stabilize the variance across replicates a two-way normalization strategy was used (see methods). Differentially expressed transcripts were identified using EdgeR. 1207 differentially expressed transcripts (>1 LogTPM, >1 log fold changes, 0.1 FDR, <0.05 pVal) were clustered into upregulated (270) and downregulated (937) *N2*^*RA/RA*^ genes (Figure 7A). Down regulated genes included the validated Notch MZB targets *Hes1, Hes5*, and *Dtx1* and upregulated genes included *FoxM1, E2F1*, and *Myb* (Figure 7B). These three genes are known to promote cell cycle, DNA replication and DNA repair in B-Cells (Figure 7C, (Lefebvre et al., 2010; Lei et al., 2005)). Indeed, all the GO term for biological process for the upregulated genes were associated with the cell cycle (Figure 7B’; Table S3), with clear enrichment for E2F1 targets (Table S3). The downregulated genes did not enrich for any significant GO terms.

**Figure 7.**
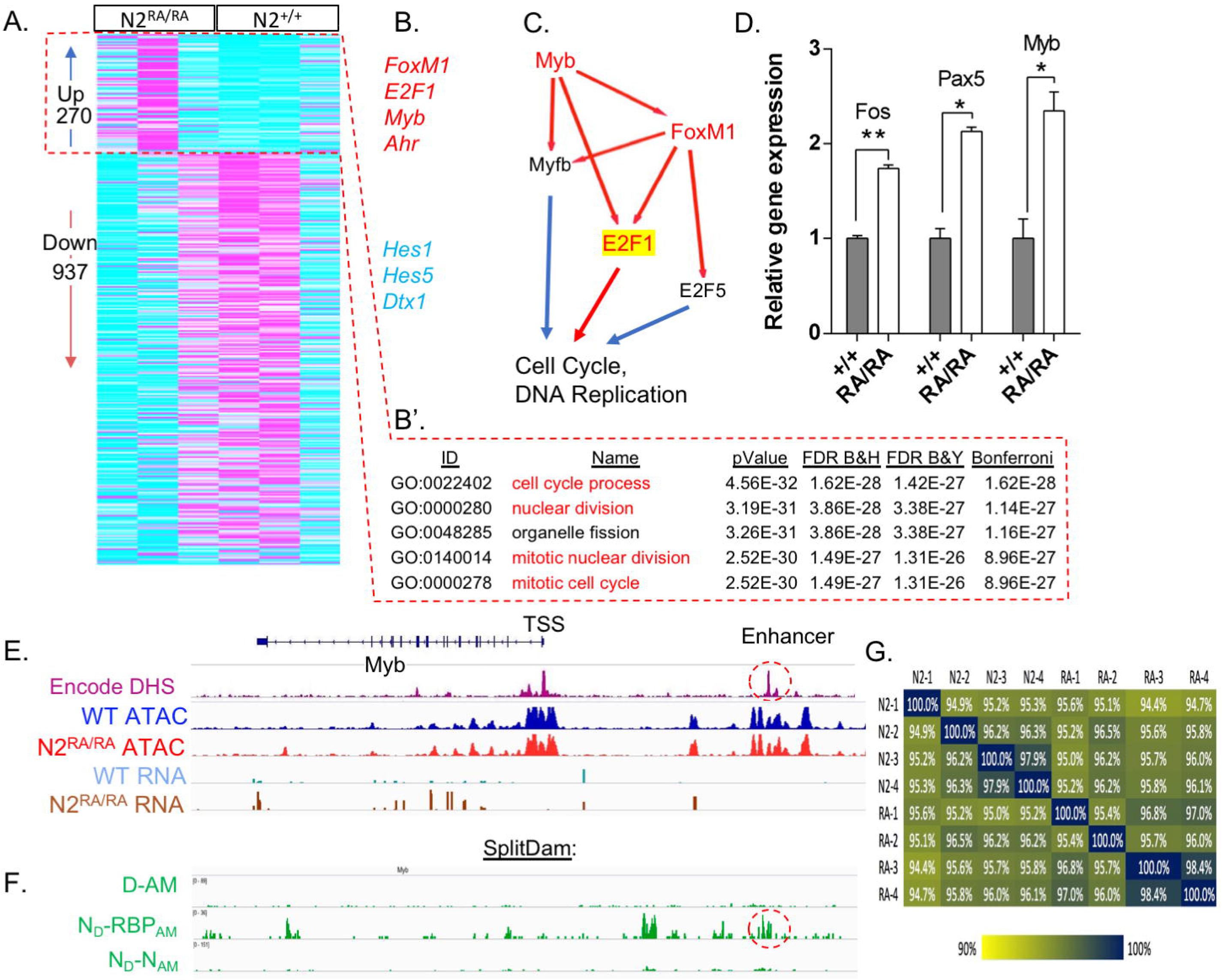
Mite infested *N2*^*RA/RA*^ activate a proliferation module in MZB. (A) RNAseq analysis identifies enrichment for genes trending down or up in *N2*^*RA/RA*^ MZB realative to wild type controls. (B) Down trending genes include classical dimer-dependent Notch targets in MZB, *Hes1* and *Hes5*, and *Dtx*. Up regulated genes include *FoxM1, Myb*, and *E2F1*. (B’) clusters 6 and 7 are highly enriched for GO terms associated with replication. (C) A schematic of the replication module controlled by *Myb*. (D) QPCR validation for *Fos, Myb*, and *Pax5*. (F) The *Myb* locus analyzed for chromatin accessibility (Encode DHS and ATAC-seq data generated in this study), transcription (RNA), and DNA methylation by DAM methyl transferase complementation (SplitDAM). Control D/AM haves methylation patterns are compared to methylation by Notch/RBPj pairs (both dimers and monomers) and Notch/Notch pairs (only dimers). See text and (Hass et al., 2015) for additional detail. (G) Spearman correlation table measures ATACseq peaks similarity matrix between all replicates. All are >92% similar.

Given that Myb can act as an inducer of FoxM1 and E2F1 (Lefebvre et al., 2010), we confirmed its transcript levels were elevated by RT-qPCR (Figure 7D). *Fos* and *Pax5* were also found to be increased within *N2*^*RA/RA*^ MZB. *Myb* could be activated by altered accessibility or by increased burst frequency and/or size, affecting amplitude. To assess accessibility, we performed ATACseq on MZB cells isolated from wild type and *N2*^*RA/RA*^ mice (4 biological replicates each). We mapped reads under the peaks and used EdgeR to identify differential accessibility across the genome between samples (see methods for detail). 89059 peaks were mapped, of which 87575 were present in both *N2*^*RA/RA*^ and wild type MZB cells with all eight samples being highly related (R>92%; Spearman Correlation; Figure 7G). Hence, genome accessibility changed minimally between these MZB cells with only 984 peaks (1.1%) enriched in *N2*^*RA/RA*^ and 500 (0.56%) in wild type. We next used GREAT to assign genes to all peaks. Based on these assignments, 14698 peaks mapped near differentially expressed (DE) genes, and of these only 108 were unique to wild type and 122 were unique to *N2*^*RA/RA*^. Importantly, none of the differently enriched peaks were present at the *Myb, FoxM1, Dtx1, Hes1* or *Hes5* loci (Supplemental Figure S7). Next, we analyzed DNAse hypersensitive peak upstream of the *Myb* locus in the ENCODE datasets, and identified an enhancer accessible in several cell types, including the kidney. The genome of a kidney-derived cell line used to overexpress constitutively active NΔE and a dimer deficient version, NΔERA (Supplemental Figure S1) was examined by SplitDamID (Hass et al., 2015). We detected strong binding to the *Myb* enhancer using Notch/RBP complementing pairs, but not by Notch dimers. This data suggested that the *Myb* enhancer responds to monomeric NTC but not to dimeric NTC. Collectively, these data suggest that the changes in MZB gene expression in the *N2*^*RA/RA*^ animals are not due to dramatic changes in chromatin accessibility but are rather due to changes in gene expression caused by the loss of Notch dimer NTC complexes at some other loci, most likely the SPS-dependent negative regulators *Hes1, Hes5*, and *Nrarp* (Hass et al., 2015; Severson et al., 2017).

## Discussion

The possibility that DNA binding site architecture contributes to Notch signaling outcomes has been considered in past studies (Liu and Posakony, 2012, 2014). NTCs can bind to enhancers with CSL binding sites as monomers that function independently of one another. In such a model, the probability of target activation will depend upon the relative abundance of nuclear NICD and the number of CSL sites at a given enhancer (Gomez-Lamarca et al., 2018; Kueh et al., 2016; Schroeter et al., 1998) and/or burst size (Falo-Sanjuan et al., 2019; Lee et al., 2019). SPSs are found in enhancer regions of up to 30% of mammalian Notch targets including *Nrarp/NRARP, Hes1/HES1, Hes5*, and *Myc* (Arnett et al. 2010, Cave et al. 2005, Liu et al. 2010, Ong et al. 2006, Severson et al. 2017, Yashiro-Ohtani et al. 2014). We have previously documented that ∼2500 SPSs were bound by N1ICD dimers in the mouse genome, and that dimerization-deficient Notch molecules bind poorly to SPSs *in vivo*, even at high NICD concentrations (Hass et al., 2015). In the murine kidney cell line, mK4, ≃15% of Notch targets required NICD dimerization. SPSs and cooperativity also proved critical for the oncogenic activity of N1ICD in murine T-cells, where NICD dimerization at a distant enhancer was required for *Myc* activation (Liu et al., 2010b). Thus, Notch targets are either agnostic to dimerization or sensitive to its presence, leading to the hypothesis that cooperativity may modulate Notch responses at physiological NICD concentrations. Recently, evidence that SPS contributed to increased probability of activation and increased duration during *Drosophila* embryonic development were reported (Falo-Sanjuan et al., 2019).

While our study concludes that embryonic development in the mouse proceeds normally in the presence of Notch receptors lacking cooperativity, it nonetheless uncovered important contributions of cooperativity to mammalian development and homeostasis under environmental stress. Notch^RA^ phenotypes are greatly enhanced by reduced gene dosage in Notch1 (causing ventricular septal defects (Donovan et al. 2002, MacGrogan et al. 2010, Sakata et al. 2002)) or in both Notch1 and Notch2 (gut). The developmental phenotype uncovered when N1^RA^ dose is reduced (in *N1*^*RA/–*^) resembles other Notch pathway deficiencies known to cause a wide range of heart defects including ventricular septal defects (VSD) (Donovan et al. 2002, MacGrogan et al. 2010, Sakata et al. 2002), and are milder than a weak hypomorphic Notch1/2 allele on the B6 background (Liu et al. 2015). Interestingly, dosage alone did not impact the ability of the *N2*^*RA*^ allele to control development and homeostasis of the dosage-sensitive marginal zone B cells in parasite-free mice, even in the hemizygote state (*N2*^*RA/–*^).

Strikingly, loss of cooperativity-dependent contributions to homeostasis becomes acute when mutant animals are burdened with exoparasites, namely fur mites. Interestingly, when the environment includes unmanaged exoparasite infestation, dimerization-deficient cooperativity mutant receptors can behave as hypomorphic (loss-of-function) alleles in some contexts (intestinal barrier formation, ventricular septum formation, intestinal stem cell self-renewal, female survival), remain indistinguishable from the wild type allele in many other tissues, or can drive the same disease as a gain-of-function allele (splenic marginal zone lymphoma, SMZL) in a cell type that otherwise seemed to be unaffected by the mutation. Note that in *N2*^*RA/RA*^ mice, Notch pathway activity in T-cells and skin, where *Notch1* is present, are identical to the wild type. This provided the rationale for the molecular analysis – the defect is more likely to reside in the B-cell linage than elsewhere, though this will remain to be examined more fully in the future. We propose these behaviors can be explained by assuming that the balance between targets agnostic to dimerization (*Myb*) and those dependent on cooperativity on SPS sites (*Hes1, Hes5, Nrarp*) varies by cell type (Figure 8). Moreover, these differences in dependence upon SPS vs non-SPS target gene regulation between tissues can be further exposed by additional stressors such as environmental insults (i.e. exoparasites) or changes in gene dose. We have proposed before that accessible SPS can act as a “sink”, holding on to NICD that otherwise will be available to regulate monomer-dependent enhancers. If true, more N2^RA^ICD may available to regulate *Myb*. Importantly, this insight may translate to other transcription factors where variants of uncertain significance are associated with developmental syndromes or neoplastic disease. Such mutations may control molecular behaviors integrating the environment with specific cooperative interactions in the affected tissues.

**Figure 8.**
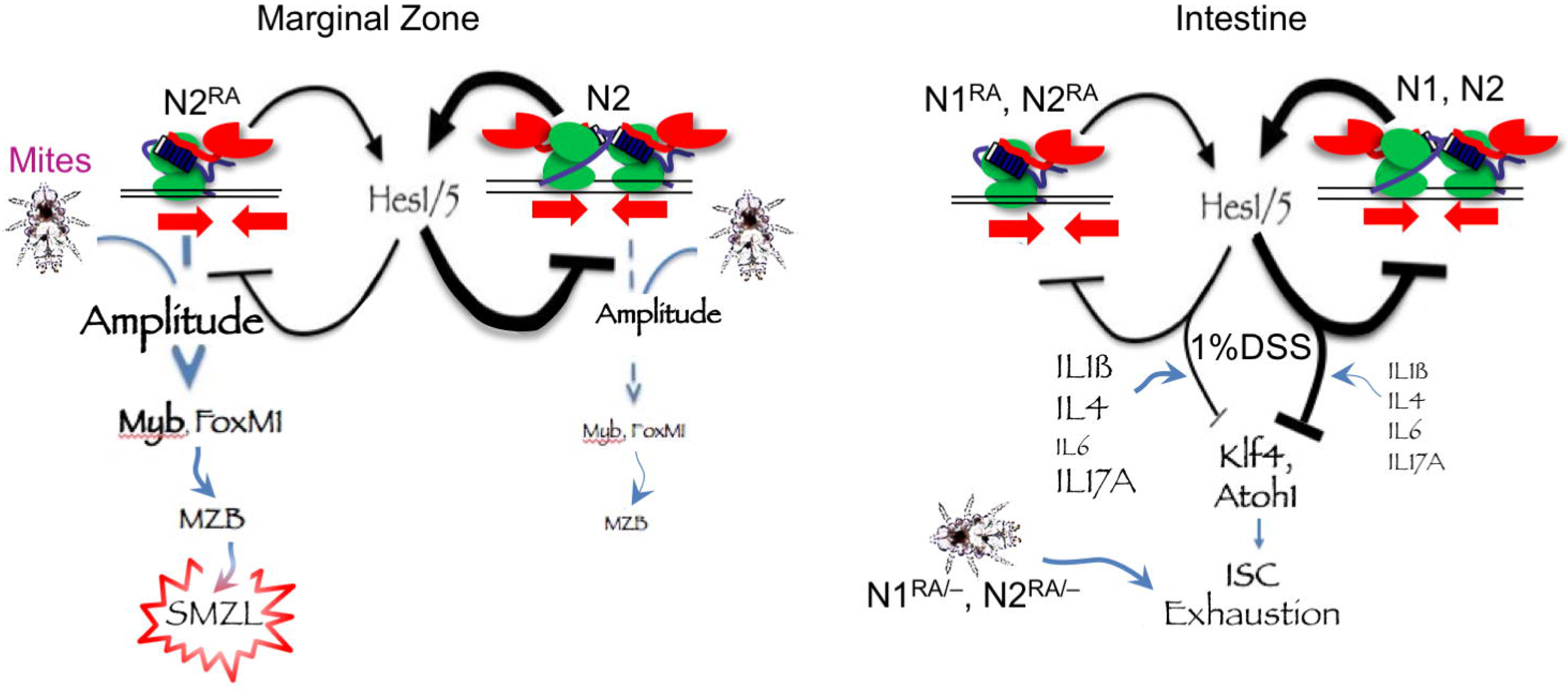
A schematic summary of the findings with a hypothetical mechanism. *Notch* integrates environmental cues (mites, cytokines) to drive proliferation in MZB (*Notch2)* and block differentiation in intestinal stem cells (ISC. *Notch*1 & 2). Activation of SPS-dependent *Hes* repressors creates a negative feedback tuning of transcription amplitude in MZB, and driving the response magnitude in ISC. In *Notch*^*RA/RA*^ mutants, SPS-dependent *Hes* gene expression is dampened, and increased availability of NICD^RA^ to monomer driven target due to reduce trapping on accessible SPS may also contribute. Repeated DSS treatment, or hemizygosity in *Notch*^*RA/RA*^ mutants, exposed insufficient blockade of pro-differentiation signals in ISC, most likely delivered via immune cells or cytokines (IL1ß, 4, or 17A) in DSS treated mice. Fur mite infestation creates an environment for runaway proliferation in MZB, and enhanced dosage effects in *Notch*^*RA/RA*^ mutants leading to complete loss of ISC postnatally even without DSS challenge.

The diverse cell types responding to a cutaneous parasite must reflect a systemic change driven by the parasite. Since we could replicate the impact on MZB proliferation in short exposures to dust mite extract, live parasite is not needed and the response is likely mediated by an immune cell (or cytokine) intermediate. Dermatitis is typically associated with both skin-produced cytokines (e.g., IL-33, -17C and TSLP), and a skewing of CD4+ T-cells into T Helper 2 (Th2) and their associated cytokines (e.g., IL-4,-5,-13, (Huang et al., 2003)). Similarly, fur mite infestation has been shown to increase the level of different cytokines (e.g. IL-6 and IL-12; (Johnston et al., 2009)), including Il-4 and -5 (Moats et al., 2016; Pochanke et al., 2006). While the exact axis linking anti-parasite immune responses and splenic marginal zone lymphoma, intestinal homeostasis, female/male ratios and heart development are yet to be identified, this study highlights the role of the environment in chronic disease, including ulcerative colitis and a smoldering malignancy such as SMZL. Combined with the significantly better outcomes in SMZL patients with activated *Notch2* alleles (Rossi et al., 2012), one has to wonder about the role of the immune environment in onset and progression of human SMZL patients with *Notch2* mutations.

## Methods

### Animals

Animals were housed at CCHMC’s animal facility in accordance with IACUC’s animal welfare regulations. The use of animals for the experiments described in this work was approved through protocol number IACUC2018-0105. To generate the RA alleles, three gRNAs (g25, g26 and g27) that target sites surrounding the R^1934^ codon in N2 were tested for their genome editing activities. Briefly, plasmids carrying the guide sequence under a U6 promoter and Cas9 were transfected into MK4 cells. Cells were harvested two days after transfection and PCR amplification, with primers flanking the target sites, was performed on genomic DNA isolated from the transfected cells. PCR products were then subjected to T7 endonuclease I (T7E1) assay. Cleavage products generated by T7E1 are indicative of indels present in the PCR products as a result of genomic DNA breakage caused by Cas9-gRNA and the subsequent repair by NHEJ. A gRNA (g3) for Tet2, which was known to have good editing activity, was included as positive control. Sixteen mice were obtained from injection of g27 (GGCATCCAGATCGGTTACA), Cas9 mRNA and a donor oligo (shown in Figure 1, g27 underlined in blue) into one-cell embryos. Digestion of PCR products with BglII showed that eight contained products with the BglII site, four of which were homozygotes. Sequencing of PCR products confirmed that all had the correct R^1934^A mutations. Since we mated the homozygote mice to each other, we identified potential second site mutations with the CCTop tool ((Stemmer et al., 2015); https://crispr.cos.uni-heidelberg.de). g27 matches to Notch2 and to a few loci with four mismatches (*Lemd1, Ccm2, Papolg, Tubb2b* and *Tubb6*) falling within an Exon and next to a PAM sequence (no loci had fewer than four mismatches to this gRNA). PCR products from potential off-target sites were sequenced and no mutations were identified in DNA isolated from any of the founders. For Notch1, a similar process led to the selection of a gRNA (CATTCGGGCATCCAGATCTG), which was injected with Cas9 mRNA and a donor oligo carrying mutations that change R^1974^ to alanine into one cell embryos. PCR products from 24 animals were digested with BglII and XbaI, respectively. Only one heterozygote founder carried the R^1974^A mutation, confirmed by sequencing. This founder was outcrossed into our mixed colony, having a single possible 4-mismatch target in the pseudogene *Vmn1r-ps6*. In the C57BL6line, the same gRNAs were used. A single heterozygote founder pair was bred within the C57BL6line.

### DSS treatment/Colitis induction

For colitis induction mice were given 1% to 2.5% Dextran sulfate sodium (DSS) in autoclaved drinking water for up to 14 days. Weight was measured every day and stool was checked for changes to monitor disease progression; in case of severe weight loss (>15%) or blood in stool mice were euthanized. For recovery mice were given normal drinking water for 14 days before starting the next cycle of DSS treatment, up to 4 cycles of injury and recovery. At the end of treatment mice were euthanized and tissue was collected for analysis.

### LPS injection/dermatitis induction

To stimulate the immune system with bacterial antigens we injected adult mice intraperitoneally twice with LPS (0.5ug/kg bodyweight, once a month) over an 8-week period. Blood was collected from the submandibular vein once per week to analyze IgM production by ELISA. The mice were euthanized after the eighth week and their spleen was harvested for histological analyses.

Mild dermatitis was induced by application of house dust mite (HDM, *Dermatophagoides farinae*) extract on the ears as described (Jin et al., 2016). In brief, the ear lobes of adult mice were gently stripped five times with surgical tape and 20 ug of HDM dissolved in autoclaved PBS was administered to each ear twice weekly for 6 weeks. Mice were euthanized and spleen harvested for histology and MZB numbers determination.

### Antibodies

A full list of antibodies is provided in the Key Resource table. The following antibodies were used for FACS analysis: Anti-mouse/human CD45R/B220, anti-CD23, anti-CD21/CD35 (CR2/CR1), anti-CD93 [AA4.1], anti-mouse IgM, anti-IgD, anti-CD45.1, anti-CD45.2, anti-CD3ε, anti-Ly-6G/Ly-6C(Gr-1), anti-CD11b, anti-TCRb and anti-CD5 were all purchased from Biolegend. Anti-Mouse CD19 was purchased from BD Biosciences. Rabbit Anti-Ki67 (Novocastra), rabbit anti-cleaved caspase 3 (Cell signaling), rabbit anti-Phospho-Histone H3 (Ser10) (Cell signaling), rat anti-Pax5 (Biolegend), Guinea pig anti-Cytokeratin 8+18 antibody and FITC rat anti-mouse/human CD45R/B220 (BioLegend) were used for fluorescence immuno-histochemistry. Anti-rabbit Alexa Fluor Cy3 (Jackson ImmunoResearch), anti-rabbit Alexa Fluor 647 (Jackson Immunoresearch), anti-rat Cy3 (Jackson Immunoresearch) and Alexa fluor anti-guinea pig 647 secondary antibodies were used for in fluorescence immuno-histochemistry. Notch2 (D76A6) XP (Cell signaling) and HRP-linked rabbit IgG (GE health care) secondary antibody were used for western blot analysis.

### Protein purification and electrophoretic mobility shift assay (EMSA)

The mouse RBPj (aa 53-474), mouse N1ICD (aa 1744-2113), mouse N1^RA^ICD (aa 1744-2113) and human SMT3-MAML1 (aa 1-280) proteins were expressed and purified from bacteria using affinity (Ni-NTA or Glutathione), ion exchange, and/or size exclusion chromatography as previously described (Friedmann et al., 2008). The purity of proteins was confirmed by SDS-PAGE with Coomassie blue staining and concentrations were determined by absorbance measurements at UV280 with calculated extinction coefficients. EMSAs were performed using native polyacrylamide gel electrophoresis as previously described (Uhl et al., 2010; Uhl et al., 2016). Fluorescent labeled probes (1.75 nM/each probe) were mixed with increasing concentrations of purified RBPj protein in 4-fold steps (from 2.5 to 640 nM) with or without the indicated purified N1ICD and MAML proteins at 2µM. Acrylamide gels were imaged using the LICOR Odyssey CLx scanner.

### EMSA quantitation

To quantitatively analyze the EMSA experiments we first extracted the gray scale intensity of each band in each lane of the EMSA gels (Supplementary Fig. S1 A-B). We then fitted the data to a 2-site binding model that takes into account cooperative binding to the 2nd site. The model calculates the binding probability assuming equilibrium binding kinetics (Michaelis-Menten) to two sites. Cooperativity is taken into account by assuming that the binding dissociation constant of a complex to the second site is divided by a cooperativity factor *C*, namely that 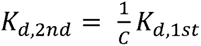. The probabilities to find the probe bound by 0,1 or 2 complexes are given by:

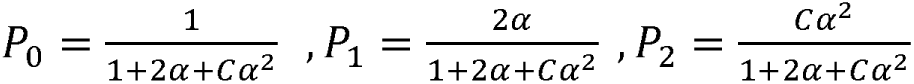

where 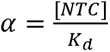 is the statistical weight associated with binding of the NTC complex to a CSL site. *K*_*d*_ is the dissociation constant to a single site. If the cooperativity factor, *C*, is equal to 1, then the binding to the two sites is non-cooperative. *C* > 1 corresponds to positive cooperativity (2^nd^ binding is enhanced). *C* < 1 corresponds to negative cooperativity (2^nd^ binding is suppressed).

Since we observe that even at high concentrations of RBPj the 1-site state is never depleted (e.g. see RBPj on CSL) we assumed that there is only some fraction of the probes, *f* < 1, that binds two complexes and another fraction, (1 − *f*), that can only bind one complex. In this case the probability to find the probe is modified to:

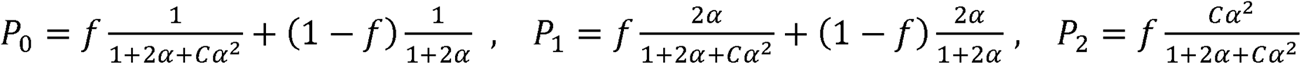

We then fit the experimental data (normalized band intensities) to these expressions. The fitting parameters are *K*_*d*_, *C*, and *f*. The parameters are extracted for each experiment separately.

To get an estimation of the confidence interval on the fitting parameters, we use a bootstrap method to randomly generate 5000 data sets with the same statistical properties as the experimental data sets (the same mean and standard deviation of band intensities). We then perform the same fitting procedure on all bootstrapped data to obtain the 95% confidence interval on the fitting parameters.

### Flow cytometry

MZB and FoB cells were sorted by FACS as described in (Chen et al. 2016). Single cell splenocytes were prepared by placing the spleen in a petri dish and mincing it with a razor blade. The disrupted spleen tissue was then transferred into a 50ml falcon tube and pipetted up and down in 5 ml of ice cold 1% BSA (Fisher Scientific) in PBS and briefly vortexed to ensure thorough disruption. Single-cell suspensions were obtained by passing the disrupted spleen tissue through a 70 um strainer (Falcon). The strainer was then rinsed with 5 ml ice cold 1% BSA/PBS and the single cell preparation pelleted by centrifuging at 2000 rpm, for 5 min followed by red blood cell lysis (BioLegend), after which the cells were pelleted at 2000 rpm, for 5 min, and resuspended in ice cold 3% BSA in PBS before cell counting. Single color antibody stained as well as unstained controls were prepared with 3 million splenocytes each, 1.5ul (0.75ug) of each antibody was added into the corresponding single-color control tube. For MZB detection, approximately 10 million cells (in 1ml 3% BSA) were distributed into Eppendorf tubes and 7.5ul of the antibody mix (2.5ul each of FITC anti-B220, PE/Cy7 anti-CD23 and PerCP/Cy5.5 anti-CD21) added into each tube. They were then incubated at 4°C for 60 minutes before sorting for B220^+^CD21^hi^CD23^lo^ MZB cells and B220^+^CD21^int^CD23^hi^ FoB cells. To isolate MZB (CD93^-^ B220^+^CD19^+^IgM^+^IgD^+^CD23^lo^), FoB (CD93^-^B220^+^CD19^+^IgM^+^CD21^+^), MZP (CD93^-^ B220^+^CD19^+^CD21^hi^IgM^hi^IgD^hi^CD23^hi^) and T2 B-cells (CD93^+^B220^+^IgM^+^CD23^+^), single cell splenocytes were incubated with the APC/Cy7 anti-B220, PE/Cy7 anti-CD93, Hoechst blue anti-CD19, FITC anti-IgM, AmCyan anti-IgD, Percp/Cy5.5 anti-CD21 and Pacific blue CD23 and FACS sorted. To analyze peritoneal B1 B-cell populations, peritoneal cells were collected as described in (Ray and Dittel, 2010) and incubated with APC/Cy7 anti-B220, PE anti-TCRb, FITC anti-CD11b, PE/Cy7 anti-CD23 and APC anti-CD5. They were then gated into B2 B-cells (B220^+^CD23^+^) and B1 B-cells (B220^+^CD23^-^). B1 B-cells were then gated into B1a (CD11b^+^CD5^+^) and B1b (CD11b^+^CD5^-^) B-cells.

For T cell analysis, thymus and spleen were harvested and disrupted by gentle grinding of tissue with a pestle (CellTreat) on a 40um cell strainer (Falcon). Red blood cells in spleen samples were lysed using ACK lysis buffer (Lonza) for 5 minutes at 4 degrees. Cells were counted and stained at 20×106 cells/ml. Cells were resuspended in PBS with Zombie Violet viability dye (Biolegend) and incubated at 4°C for 20 minutes. Cells were washed, and then resuspended in an antibody cocktail in PBS supplemented with 2% FBS (Hyclone) in the presence of 5% 2.4G2 Fc blocking antibody (in house), and incubated for 30 minutes at 4 degrees. Cells were washed, resuspended in PBS with 2% FBS, and analyzed using an LSRII (BD). Data were analyzed using FlowJo software v9.7 and average population percentages and absolute numbers were graphed using GraphPad Prism (GraphPad Software, Inc.).

### MZB and FOB culture and ELISA

MZB and FoB cells were cultured in RPMI-1640 (Sigma-Aldrich) containing 10% FBS (Sigma-Aldrich), 10µg/ml gentamicin (Gibco B.R.L) and 50µM β-mercaptoethanol (Sigma-Aldrich). 10^5^ MZB and FoB cells were seeded into U-bottom 96-well plates (Thermo Scientific) in 200µL of media. Differentiation was stimulated by adding lipopolysaccharides from *Salmonella enterica* serotype typhimurium -LPS (Sigma-Aldrich) into each well at a final concentration of 2µg/ml. The LPS-stimulated cells were cultured at 37°C, 5% CO_2_, for 5 days. To measure IgM levels by ELISA, MaxiSorp flat-bottom 96-well plates (Nunc) were coated with 200µL of 5µg/ml goat anti-mouse IgM (Southern Biotech) in 0.05% PBST (PBS-tween20) at 4°C, overnight. The plates were then washed thrice with 0.05% PBST and blocked with 3% BSA (Fisher Scientific) in PBS for 30 minutes at room temperature. Purified mouse IgM (Southern Biotech) was used to prepare a control standard. 150µl of supernatant collected from the stimulated B-cells at the end of 5-day incubation was diluted in 0.05% PBST (PBS containing 0.05% Triton-100), and along with the standard, was loaded on to the MaxiSorp flat bottom (Thermo Fisher Scientific) ELISA plates for 1hr incubation at room temperature followed by three washes with PBST. HRP goat anti-mouse IgM (Southern Biotech), diluted at 1:5000 in PBST, was added and incubated for 1hr at room temperature followed by three washes with PBST. TMB substrate solution (Thermo Fisher Scientific) was then added and developed for 10 minutes. Development was stopped with 0.16M sulfuric acid before colorimetric absorption analysis at 450nm.

### Histology

Mouse tissues were fixed overnight in 4% PFA and embedded in paraffin. Adult intestinal tissue was divided into sections corresponding to the jejunum, duodenum, ileum and colon. Each section was then folded into a swiss-roll (Moolenbeek and Ruitenberg, 1981) and then fixed, mounted and sectioned for histological analysis. For P0 animals, whole intestinal tissue was collected and then fixed and mounted whole before sectioning for histological or immunofluorescence analysis. 5µm thick sections were deposited on Superfrost plus microscope slides (Fisher Scientific), deparaffinized with xylene (Fisher Scientific), and rehydrated. Sections were stained with hematoxylin for 2.5 minutes and Eosin for 1 minute before dehydration, clearing and mounting. For alcian blue staining, sections were rinsed in 3% acetic acid (Fisher Scientific) for 3 minutes and incubated at room temperature for 30 minutes in 1% Alcian Blue 8GX (Sigma-Aldrich) in 3% acetic acid solution. The sections were then rinsed for 3 minutes in 3% acetic acid followed by 10 minutes in running water. After counterstaining with Nuclear Fast Red (Vector) for 5 minutes, slides were rinsed for 10-minutes in running water prior to dehydration, clearing and mounting. For each sample, images of at least 10 random fields of view were acquired on a Nikon 90i Upright widefield microscope. Where indicated, quantitative analysis of images was done using Nikon’s NIS-Elements Advanced Research software (NIS-AR). In order to automatically identify and quantify Alcian Blue positive goblet cells, an inhouse NIS-AR plugin was applied to identify and mask the discrete, large, dark blue Alcian Blue spots scattered on the pink-white Nuclear Fast Red counterstain (Figure 3). We then manually traced the villi in the region of interest (ROI) to exclude the Alcian Blue stained mucus. The plugin then counted the Alcian Blue positive foci within the ROI. Goblet cell numbers were determined for all acquired fields of view for each animal analyzed. The average number of goblet cells per field of view was obtained for each analyzed animal. Where multiple animals of the same genotype were analyzed, the average goblet cell number per field of view was obtained by averaging the goblet cell numbers in all fields of view from all the animals of that genotype.

### Fluorescent Immunohistochemistry

Paraffin embedded sections were deparaffinized with xylene, rehydrated, and subjected to Heat-Induced Epitope Retrieval (HIER) by boiling in Trilogy (Cell Marque) for 30 minutes. After cooling, the sections were rinsed with distilled water for five minutes followed by PBS with 0.3% Triton-X100 (Fisher Scientific) for five minutes. The sections were then incubated at room temperature for two hours in blocking solution (10% normal donkey serum (Jackson ImmunoResearch) in PBS-Triton), followed by overnight incubation with respective primary antibodies at 4°C. The sections were washed 3×10 minutes with PBS-Triton and incubated for two hours at room temperature with Fluorophore-labeled secondary antibodies in blocking solution. Unbound secondary antibody was washed three times for 10 minutes each in PBS-Triton and where indicated, counterstained with DAPI before mounting with Prolong Gold Antifade (Invitrogen). Confocal imaging was performed on a Nikon Ti-E Inverted Microscope.

### Cytoplasmic/nuclear separation and Western Blot

N2ICD stability was determined in FACS-isolated MZB and FoB cells. Sorted cells were pelleted for 5 min with 2000 rpm at 4°C and resuspended in 500 ul hypotonic buffer (HB; 20mM Tris-HCl, pH7.5, 10mM NaCl, 3 mM MgCl_2_) with added protease inhibitor (Roche). After 30min incubation on ice, 25 ul of 10 NP-40 was added and samples vortexed for 10 seconds. Nuclei were pelleted at 3000 rpm/10min at 4°C and the cytoplasmic fraction transferred into a fresh tube. The nuclei pellets were washed twice in 500ul HB and then lysed in 20ul 2X Laemmli sample buffer (120mM Tris-HCl, pH6.8, 20% glycerol, 4% SDS). The cytoplasmic fractions were concentrated using AmiconUltra – 0.5mL – 30K centrifugal filters (Merck Millipore) and mixed 1:1 with 2X Laemmli sample buffer. The extracts were separated on 6% polyacrylamide (Biorad) gel and transferred onto nitrocellulose membranes in Tris/glycine transfer buffer. Membranes were blocked with blocking solution (5% milk in 0.1% PBS-Tween20 (Fisher Scientific) for 1 hour, room temperature, and incubated overnight at 4°C with primary antibody in blocking solution. Membranes were then washed three times for five minutes with 0.1% PBS-Tween20 and incubated for 1 hour at room temperature with anti-rabbit HRP secondary antibody (GE healthcare) in blocking solution. Membranes were developed with a Supersignal West femto chemiluminescent substrate kit (Fisher scientific) and developed using Chemidoc (Biorad) detection system. Signal intensities were quantified using Image Lab (Biorad) software.

### qPCR

RNA from tissue and cells was extracted using the PureLink™ RNA Mini kit (Invitrogen) and cDNA was synthesized with SuperScript® II reverse transcriptase (Invitrogen) following the manufacturers instruction. Quantitative PCRs were performed using iTaq Universal SYBR® Green Supermix (BioRad) on the StepOnePlus™ RT PCR system. Data were analyzed using the Delta-Delta-CT methods. A full list of oligos is provided in In Key Resources Table.

### RNAseq

Total RNA was isolated from FACS-sorted MZB cells using a Qiagen MicroRNAeasy kit according to manufacturer’s instructions. RNA-SEQ libraries were generated using the Nugen Ovation RNA-Seq System V2 and 75 b paired-end reads were obtained on an Illumina Hi-Seq 2500 machine with read depths of ∼38 million following to Illumina protocols. Quality checked RNA-seq reads were mapped to mouse genome (mm10) using RSEM (v1.2.19) and quantified. Raw counts and transcripts per million (TPM) units were reported for all genes. Principal component analysis (PCA) was performed using R package prcomp. ∼60% variance was accounted for by PC 1 and 2. Based on the PCA, analysis, we excluded biological replicate 3 in both WT MZB and RA MZB from further analysis (See Table S3). Differential Expression was performed using R package RUVSeq, which normalizes and contains EdgeR. A two-way normalization strategy was used to stabilize the variance across replicates for each transcript: an upper quantile normalization followed by an empirical gene normalization using ∼5000 least differentially expressed transcripts. Two-way normalized counts were then used to perform differential expression with EdgeR. Upregulated and downregulated genes were filtered (see Table S3) and loaded to the Toppgene suite (https://toppgene.cchmc.org/enrichment.jsp) to identify significant enrichment.

### ATACseq

For sample library preparation we followed the Omni-ATAC method outlined by (Buenrostro et al., 2015; Corces et al., 2017) and purified Tn5 was generated as described (Picelli et al., 2014). Briefly, 50,000 nuclei from FACS-sorted MZB cells were processed for Tn5 transposase-mediated tagmentation and adaptor incorporation at sites of accessible chromatin. FACS isolated MZB cells were pelleted and washed with ice-cold PBS. The pellet was resuspended in ATAC-Resuspension Buffer (10mM Tris-HCl pH 7.4, 10mM NaCl, 3mM MgCl2 0.1% NP40, 0.1% Tween-20, 0.01% Digitonin) and incubated on ice for 3 minutes. The lysed cells were washed in ATAC-Wash Buffer (10mM Tris-HCl pH 7.4, 10mM NaCl, 3mM MgCl_2_, 0.1% Tween-20), inverted three times and the nuclei pelleted. The nuclei were resuspended in ATAC transposition mix (10mM Tris-HCl pH 7.6, 10mM MgCl_2_, 20% Formamide, 100nM Tn5 transposase) and incubated at 37°C for 30 minutes in a thermomixer at 1000 RPM. Following tagmentation, the DNA fragments were purified using a Zymo DNA Clean and Concentrator Kit and library amplification was performed using customized Nextera PCR primer Ad1 in combination with any of Ad2.1 through Ad2.12 barcoded primers as described (Buenrostro et al., 2015). The quality of the purified DNA library was performed utilizing an Agilent Bioanalyzer 2100 using High Sensitivity DNA Chips (Agilent Technologies Inc., Santa Clara, CA). The samples were pooled at a concentration of 5nM and run on an Illumina HI-SEQ 2500 sequencer (Illumina, Inc. San Diego, CA) to obtain paired-end reads of 75 bases (PE75). ATAC sequencing was carried out on two conditions MZB wild-type and RA. For each condition, 4 biological replicates were sequenced, in single end fashion. Read depth for replicates varied from ∼ 50 to 98 million reads thereby giving us average read depth of ∼67 million reads. Quality check, mapping and peak calling was performed using CSBB-v3.0 [Process-ChIP-ATAC_SingleEnd]. CSBB uses fastqc, bowtie2 and macs2 [parameters: --nomodel --shift 37 -- extsize 73] respectively. Duplicate mapped reads were removed before peak calling. Mapped reads in bam format was converted to bigwigs using deeptools (<u>deeptools.ie-freiburg.mpg.de</u>) for visualization purposes. For assessing open nucleosome region differences from MZB WT to RA MZB, we performed differential peak analysis. Within each sample type only peaks with 75% replication and at least 50% overlap among biological replicates were used for differential peak analysis. Further, peak sets passing above defined criteria were merged (at least 1bp overlap) using bedtools merge and then number of reads mapping under each peak for each replicate was inferred using FeatureCounts program. Finally, differential peak analysis was performed using EdgeR. No statistically significant changes in peaks (ATAC regions) between WT MZB and RA MZB were identified.

## Key Resources

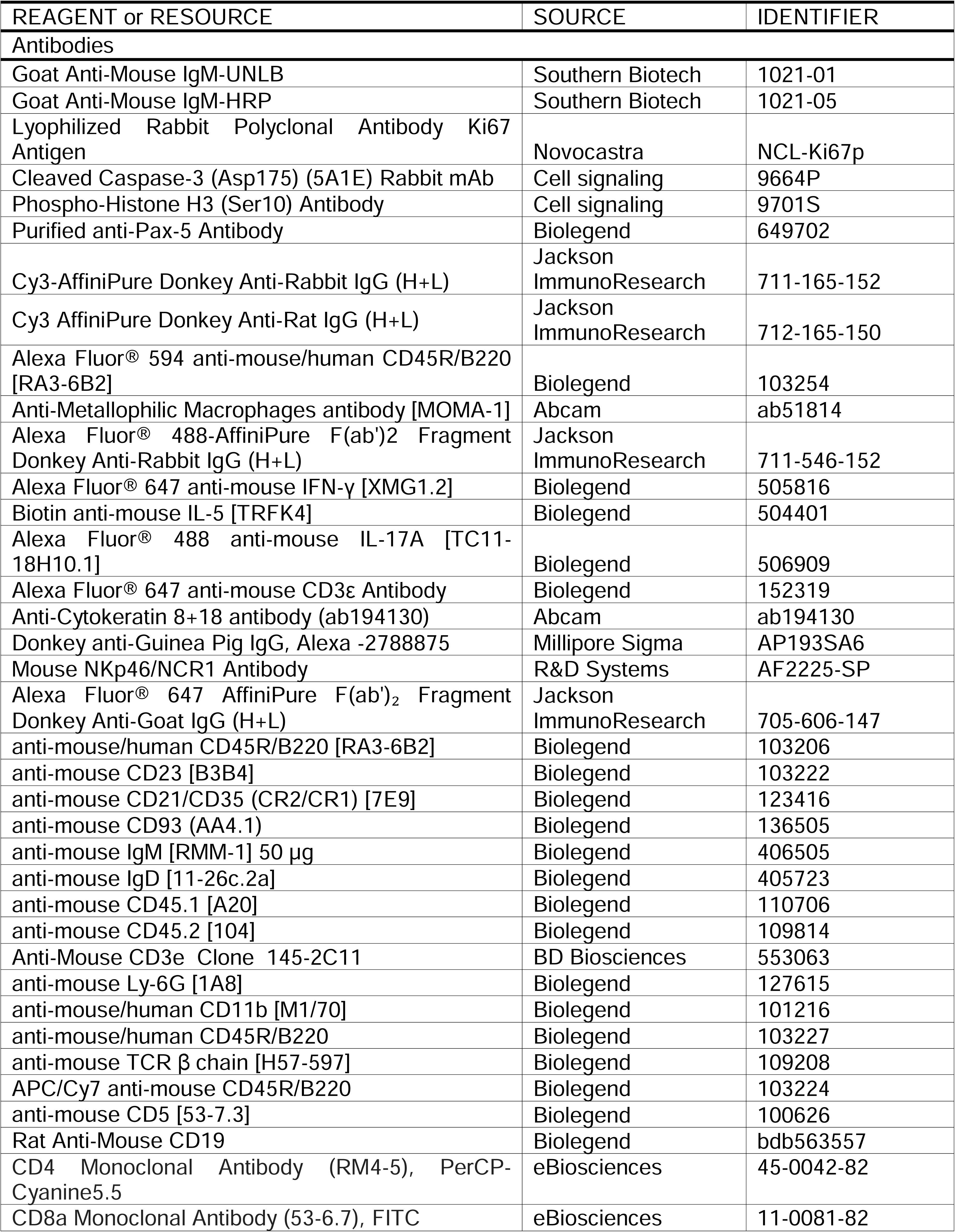
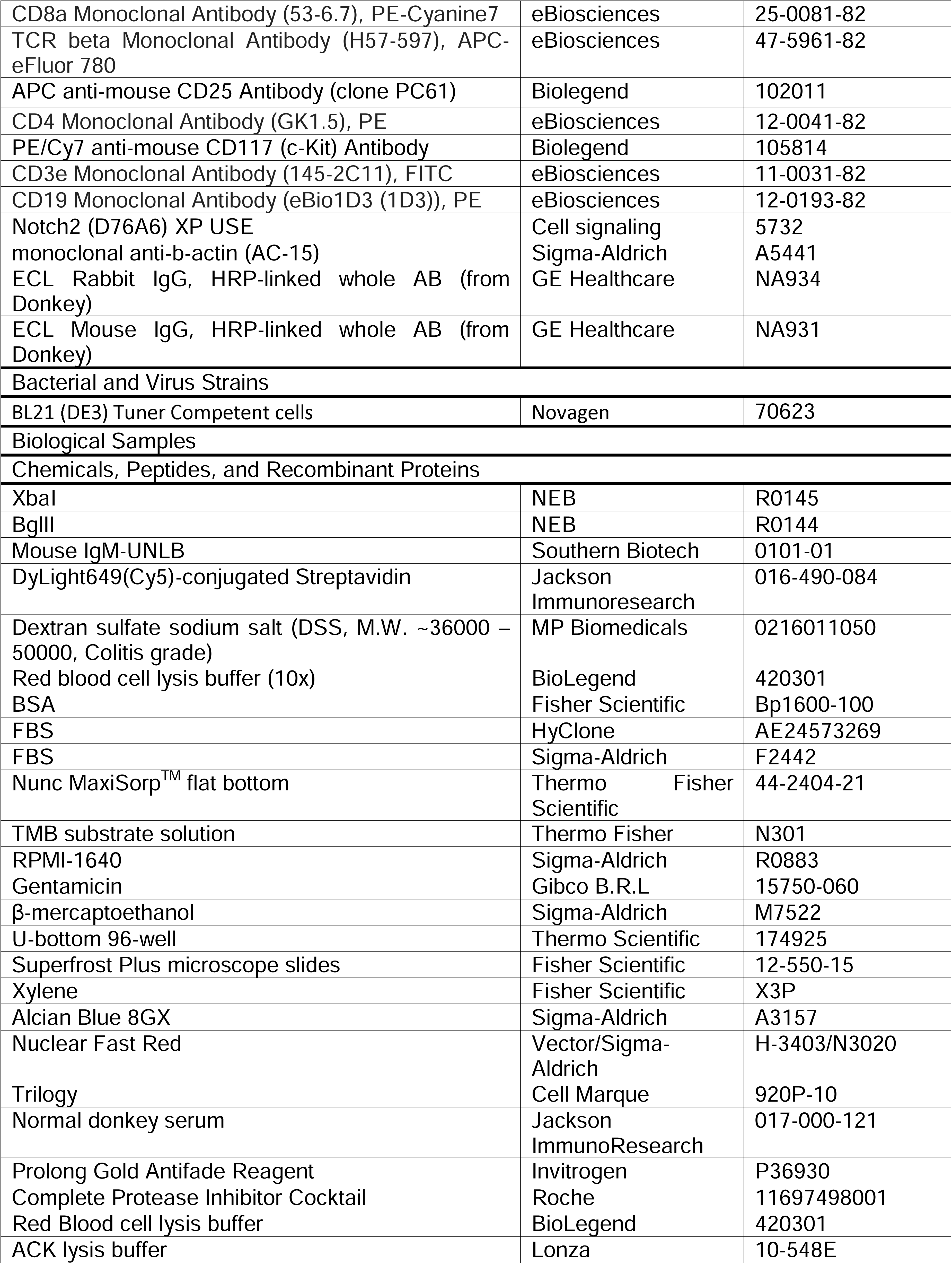
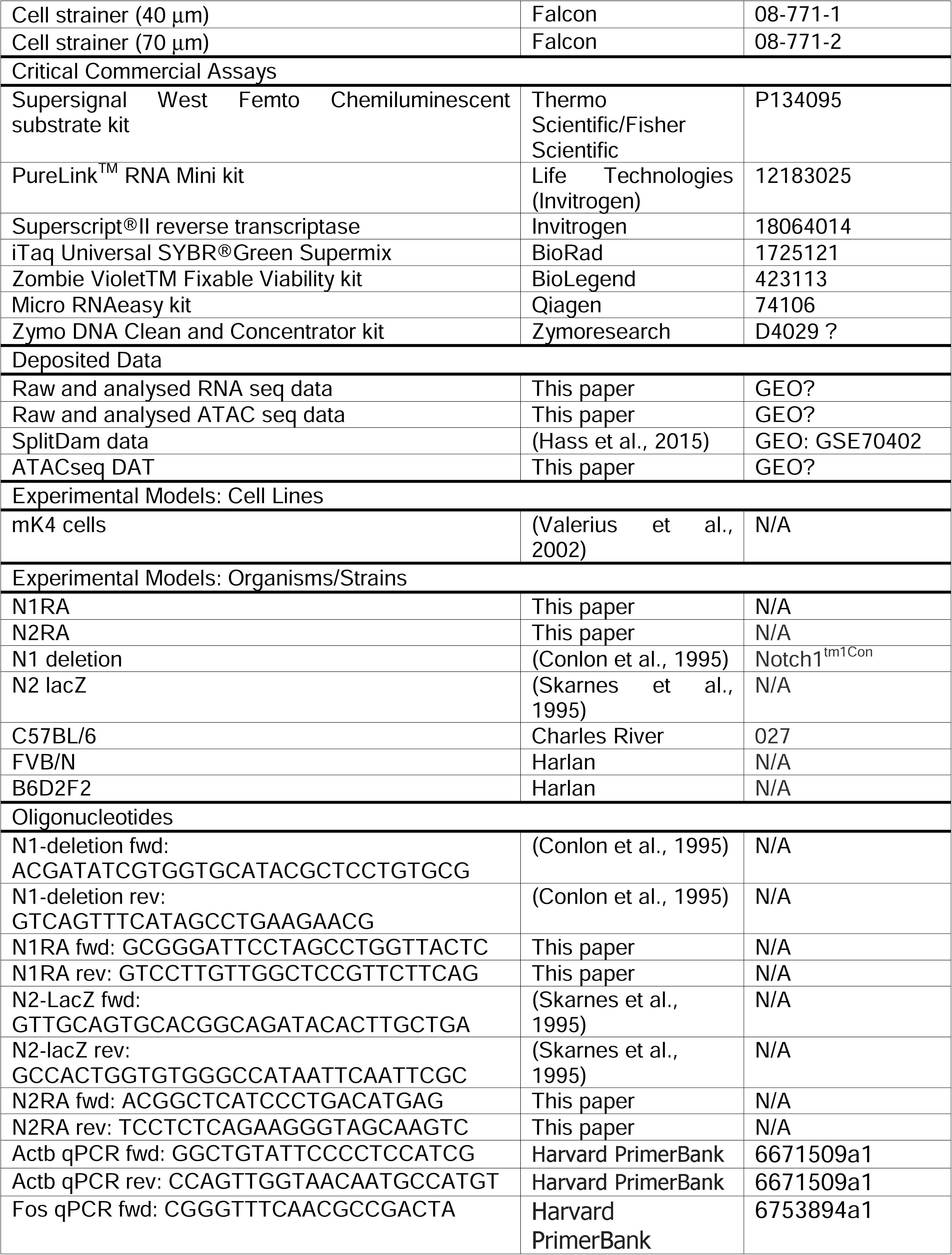
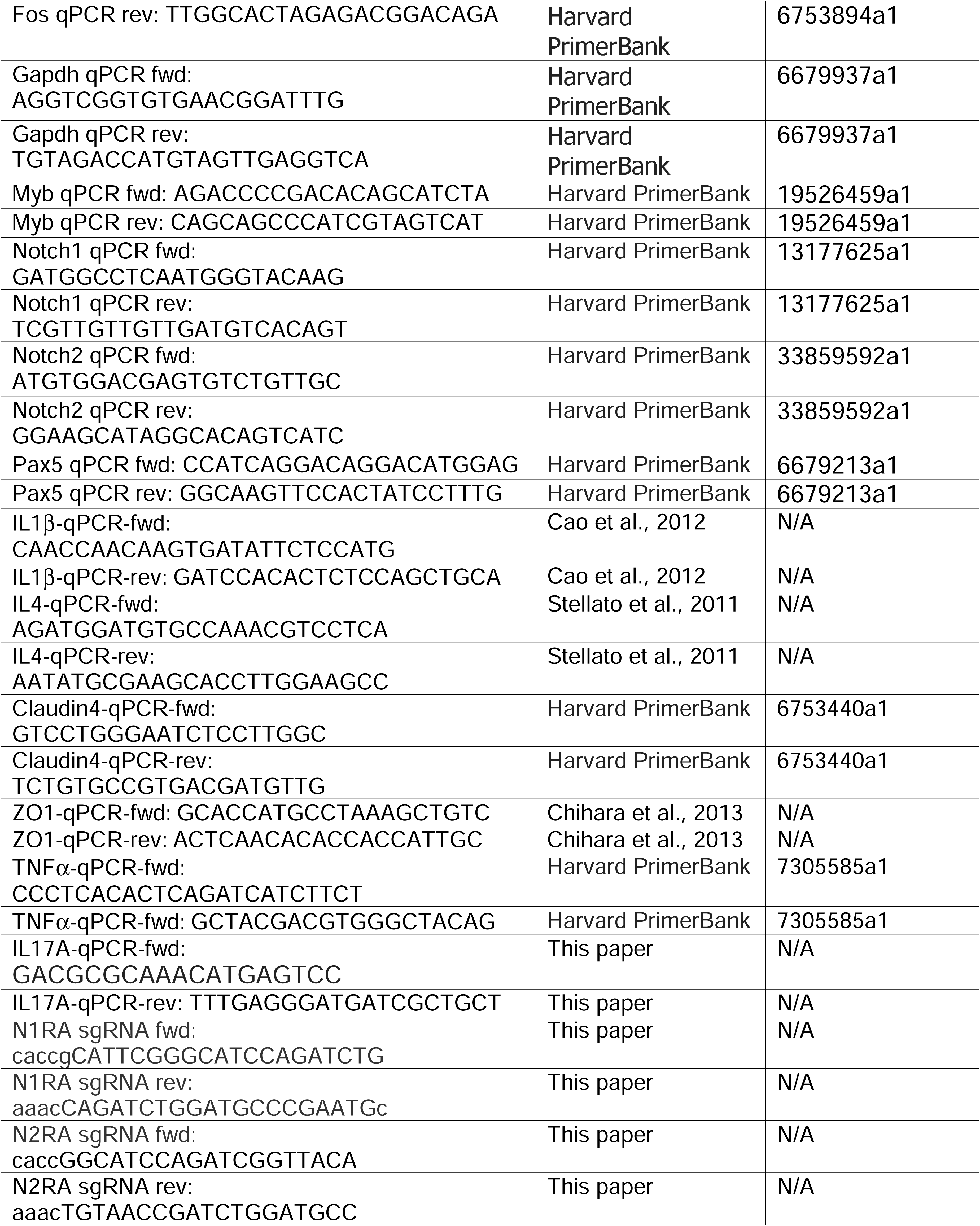
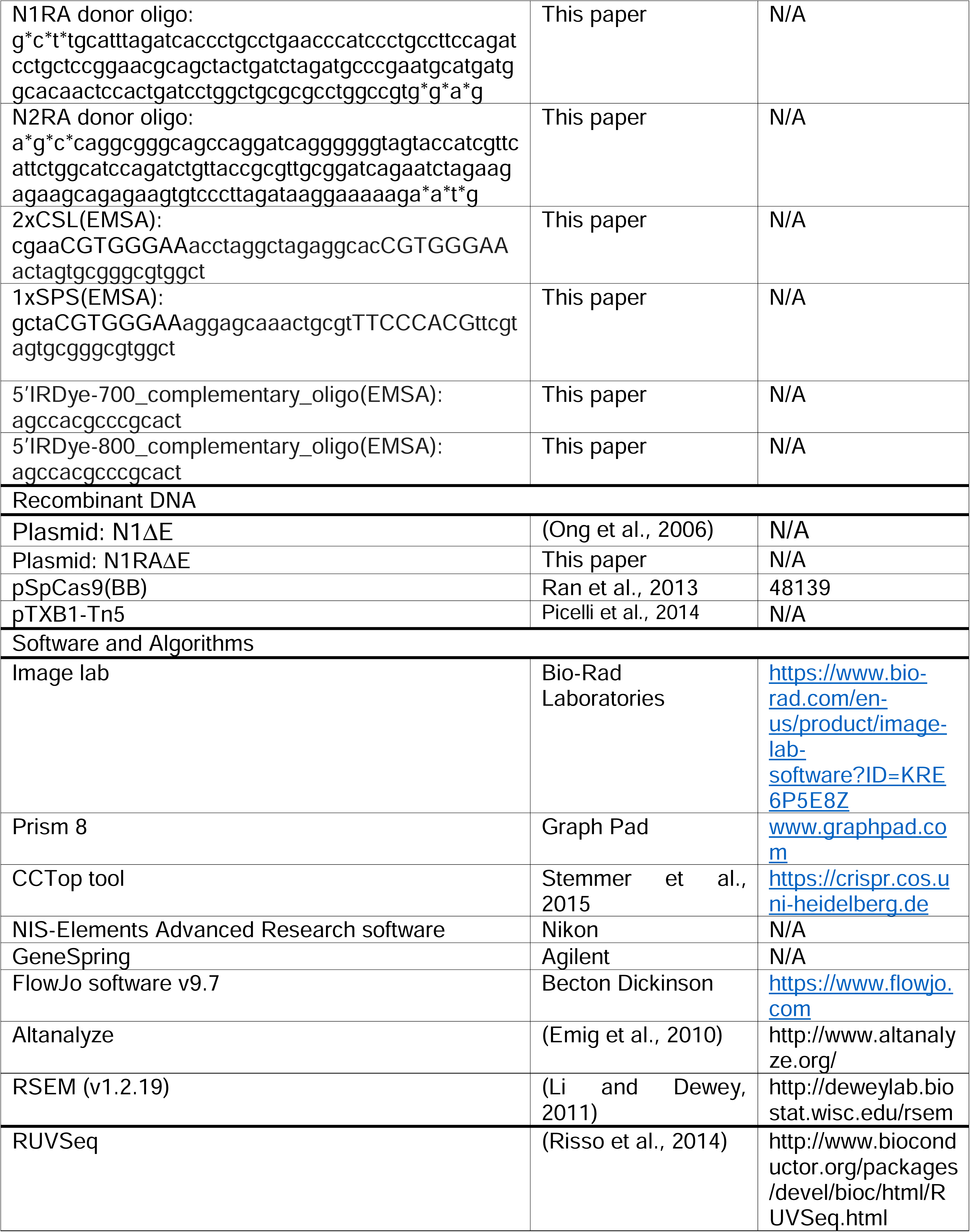
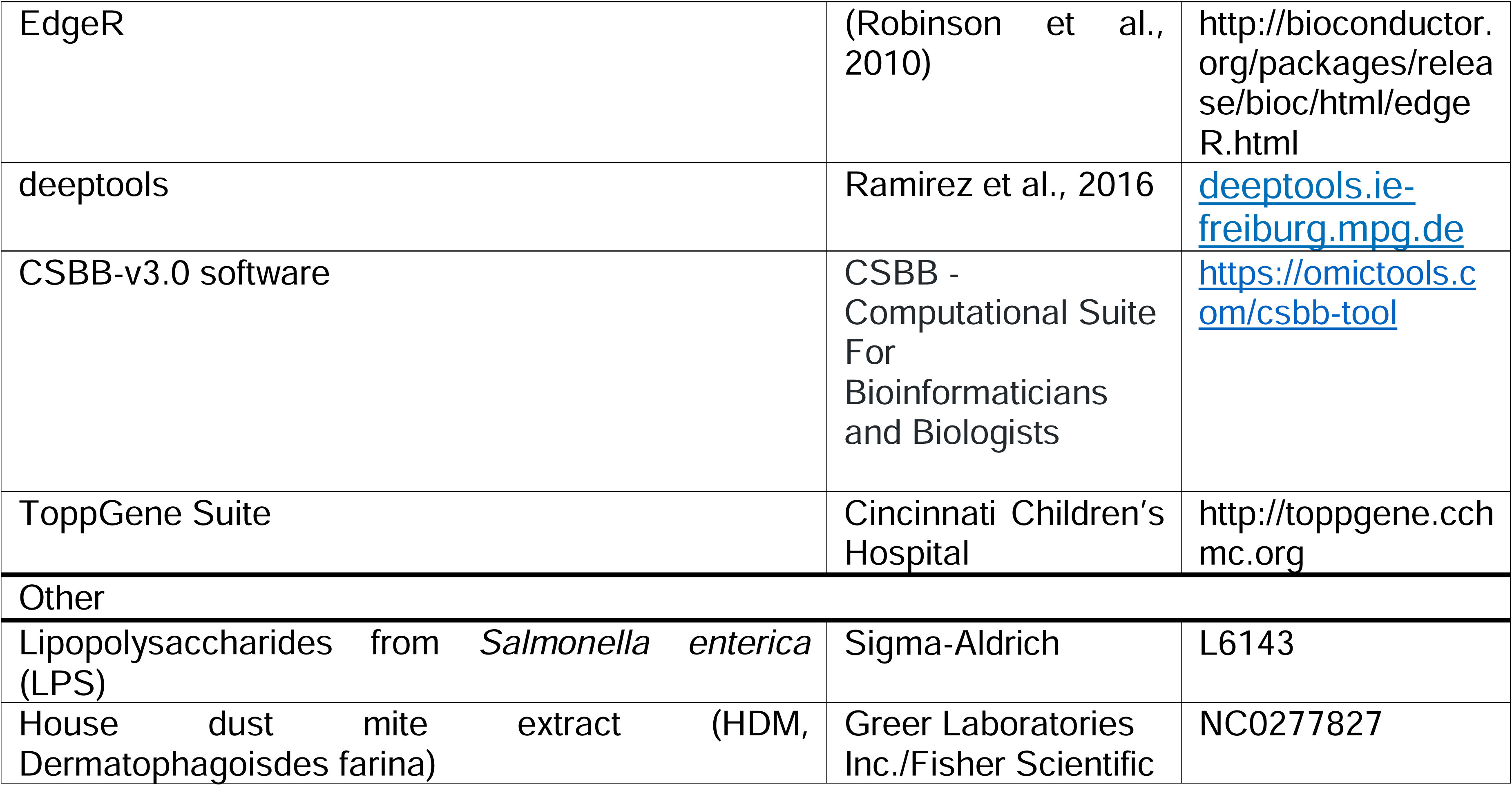

## Acknowledgments

The authors wish to thank Dr. Matt Kofron with help with imaging, Dr. Yueh-Chiang Hu and the genome editing core for generating the NRA alleles, and members of the Kopan lab for comments and encouragement during the study. Dr. Stephan Waggoner, Kelli VanDussen and members of the DB and Notch community for valuable advice. Many thanks to Dr Sai Tammula and the veterinarians at CCHMC. Cell sorting was supported by NIH S10OD023410 and 01DK106225 to the CCHMC Research Flow Cytometry Core. BG and DS were supported at a grant from the National Science Foundation and the Binational Science Foundation (NSF/BSF - 1715822). SJS was supported by NIH T32CA009140 and the American Cancer Society PF-15-065-01-TBG. WSP was supported by NIH R01 CA215518 (WSP). RK, FMK, KP, NW, and EB were supported by MIH RO1 CA163353 (RK). RK and QD were supported by the William K. Schubert Endowed Chair at Cincinnati Children’s Hospital Medical Center. QD receives a small stipend from the Chinese government.

## Authors contributions

FMK generated most of the data in mite infested mice and contributed to writing of the manuscript; KP contributed most of the data in the mite-free colony and supervised QD during the DSS experiments. NW contributed the initial establishment of the MZB phenotype. RAK/ZY purified the proteins, YK, NE, DS and BG analyzed cooperativity, with YK/BG performing EMSA experiments and NE/DS calculating the cooperativity factor. SJS and WSP analyzed *N1*^*RA*^ T-cells. EB performed the RNA and chromatin analyses and supervised NW. BA performed the RNAseq clustering and enrichment analysis. EB/RK conceptualized the study and analyzed the data with input from KP, FMK, and QD. RK wrote the manuscript. All authors read and edited the final version.

