## Supplemental Information for "Exposure to Mites Sensitizes Intestinal Stem Cell Maintenance, Splenic Marginal Zone B Cell Homeostasis, And Heart Development to Notch Dosage and Cooperativity" (from another version of this preprint: version 2, 2020-02-27)

**Supplementary figure legends**

**Supplementary Figure S1: Loss of Notch1 dimerization suppresses target gene expression in MK4 cells (supporting Figure 1).** (A-B) EMSA data for purified proteins binding to probes containing two CSL (A) or SPS (B) sites. The number of balls marks the occupancy of sites, arrows below indicate orientation of binding sites in probes. (C-D) The average sites filled were plotted against RBPj concentrations to calculate the cooperativity factor “C” on CSL (C) and SPS (D) probes. (E) Relative expression of Notch target genes in mK4 cells overexpressing either N1∆E or N1^RA^∆E as determined by RNA Seq (all replicates are shown, values ranked per row. Green arrows indicate targets elevated more in N1^RA^∆E, red arrows indicate targets repressed in in N1^RA^∆E relative to control.

**Supplementary Figure S2: Normal development of T cell compartments in *N1^RA/RA^* mice (supporting Figure 1).** Thymi and spleens were isolated from *wt*, *N1^+/RA^*, or *N1^RA/RA^* mice, and the T cell compartment was analyzed. (**A**) The absolute number of thymic single- and double-positive T cells (left) and developing T cells (right) was assessed by flow cytometry. (**A’**) The average number of cells in the thymus (left) and average thymic weight (right) are shown. (**B**) The percentage (left) and absolute number (right) of T cell subsets in the spleen were assessed by flow cytometry. (**B’**) The average number of splenocytes (left) and average spleen weight (right) are shown. (n=3-6 mice per genotype; error bars = +/- SEM)

**Supplementary Figure S3: Proliferation potential in wild type vs mutant intestines with or without fur mite exposure (supporting Figure 4).** Sections of P0 and P1 intestines from animals born to fur mite infested dams (**A-H’**) and born after fur mite eradication (**I-J’**) were stained for Ki67 (**A-D’, I** and **I’**) and phosph-H3 (**E-H’, J** and **J’**) to assess for proliferation in the crypts. *N1^+/-^;N2^+/-^* are indistinguishable from *wt* (**A-B’, E-F’**), while proliferation is significantly reduced in *N1^+/-^;N2^RA/RA^* and *N1^RA^/-;N2^RA/-^* crypts (**C-D’, G-H’**). Proliferation in *N1^RA/-^;N2^RA/-^* is still decreased compared to heterozygous littermates after fur mite eradication (**I-J’**).

**Supplemental Figure S4: Like wildtype MZB cells, *N2^RA/RA^* MZB cells have a robust proliferative response upon LPS stimulation and *N2^RA/RA^* spleen display germinative centers (supporting Figure 5).** Isolated MZB from *N2^RA/RA^* and *wt* littermates were cultured and stimulated with LPS. Both genotypes proliferate after stimulation (A). Sections of *N2^RA/RA^* and *wt* spleens were stained with Ki67 and phosphor-H3 to detect proliferation and Caspase3 for apoptosis (**B**). Proliferation in germinative centers were detected in dimer deficient mice, but not in *wt*. Apoptosis was not altered.

**Supplementary Figure S5: Spleen size as a function of age In the absence of pathogens; *N1^+/RA^;N2^RA/RA^* enlarged spleens and lymph nodes of mite infested mice have a high proliferative and mitotic index.** (**A**). In the absence of fur mites, an increase in spleen size with aging was observed in *N2^RA/RA^* mice (RA), but not in mice with other genotypes (+) housed in the same colony. (**B, D**). The enlarged spleens and lymph nodes from aged *N1^+/RA^;N2^RA/RA^* mice showed increased staining for Ki67 and phosphor-H3 indicating proliferation Apoptosis was slightly increased in enlarged spleens as shown by Caspase3 stain (**C**). (**E**) Low magnification of spleens from aged *N2^RA/RA^* and *N1^+/RA^;N2^RA/RA^* mice infested with fur mites show expansion of white pulp.

**Supplementary Figure S6: Dimerization of NICD does not affect protein stability.** Western blot analysis of nuclear (N) and cytoplasmic (C) preparations of sorted MZB shows no difference in N2 stability between *wt* and *N2^RA/RA^* mice (**A**); immunoblotting of α-tubulin and histone-h3 confirms separation of cytoplasmic and nuclear fractions, respectively. Quantification of the nuclear N2ICD relative to cytoplasmic total N2 reveals no difference between wt and dimer deficient N2ICD stability (**B**).

**Supplemental Figure S7: Accessibility at dimer responsive loci is not altered by dimer deficiency.** ATAC-Seq data reveal that DNA accessibility was not altered at dimer responsive loci with upregulated (**A**) or downregulated expression (**B**), compared to loci with dimer independent expression (**C**); *wt* in blue, *N2^RA/RA^* in red. Expression of the genes was changed as shown by RNA-Seq data (arrows indicate up- or downregulation); *wt* in cyan, *N2^RA/RA^* in brown.

**Supplementary Figure S1**
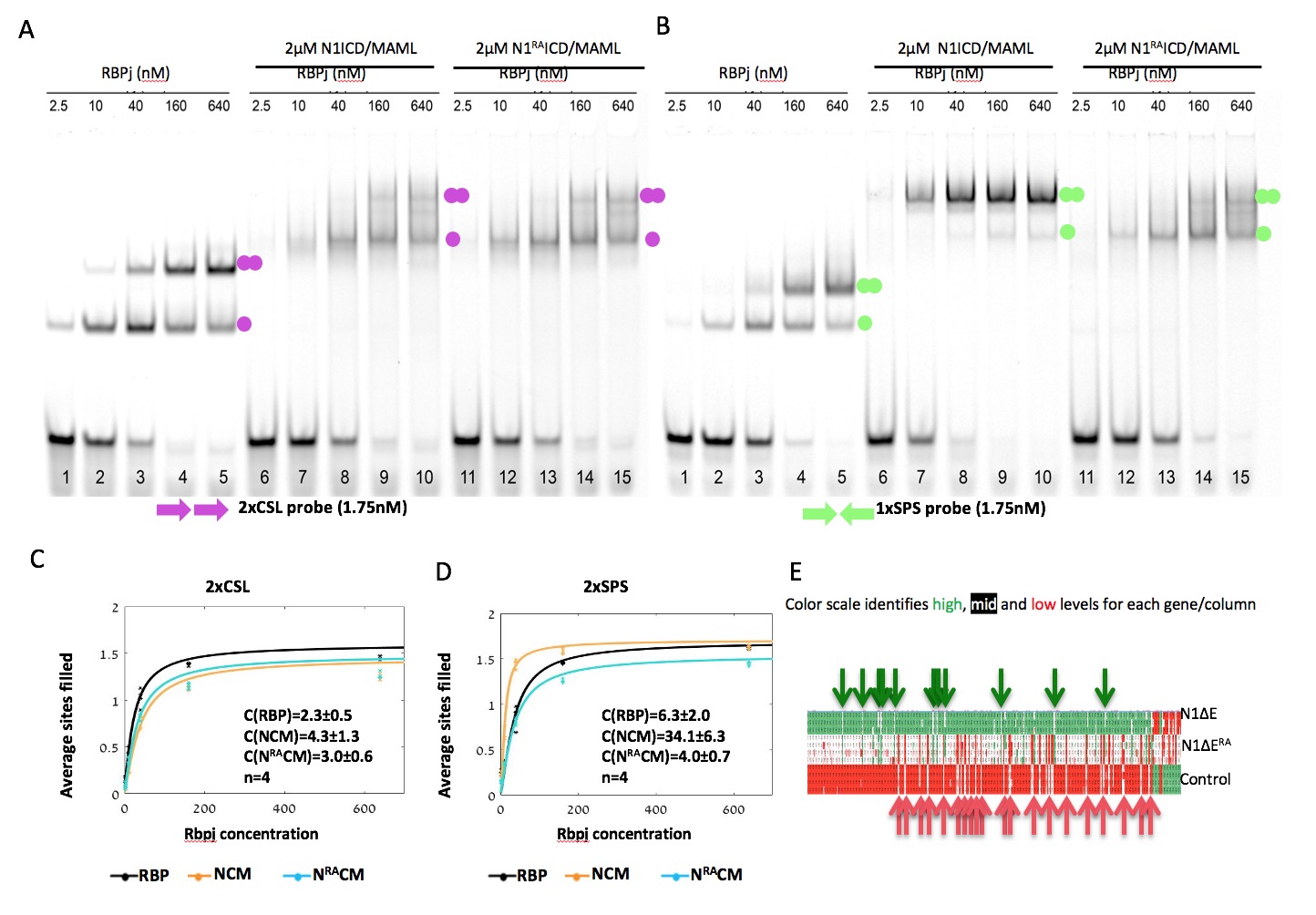
**:**

**Supplementary Figure S2:**

**Supplementary Figure S**
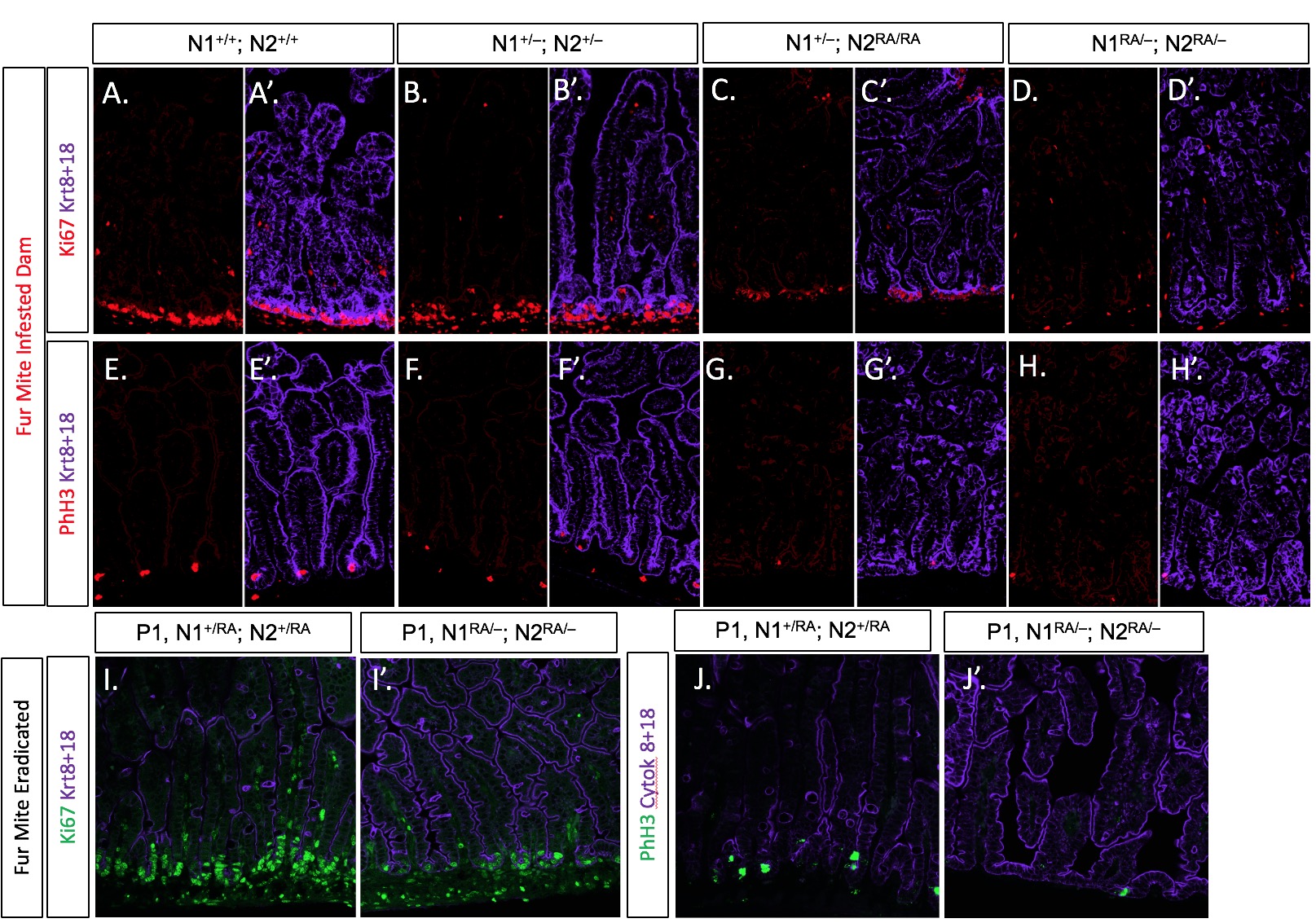
**3:**

**Supplementary Figure S4:**
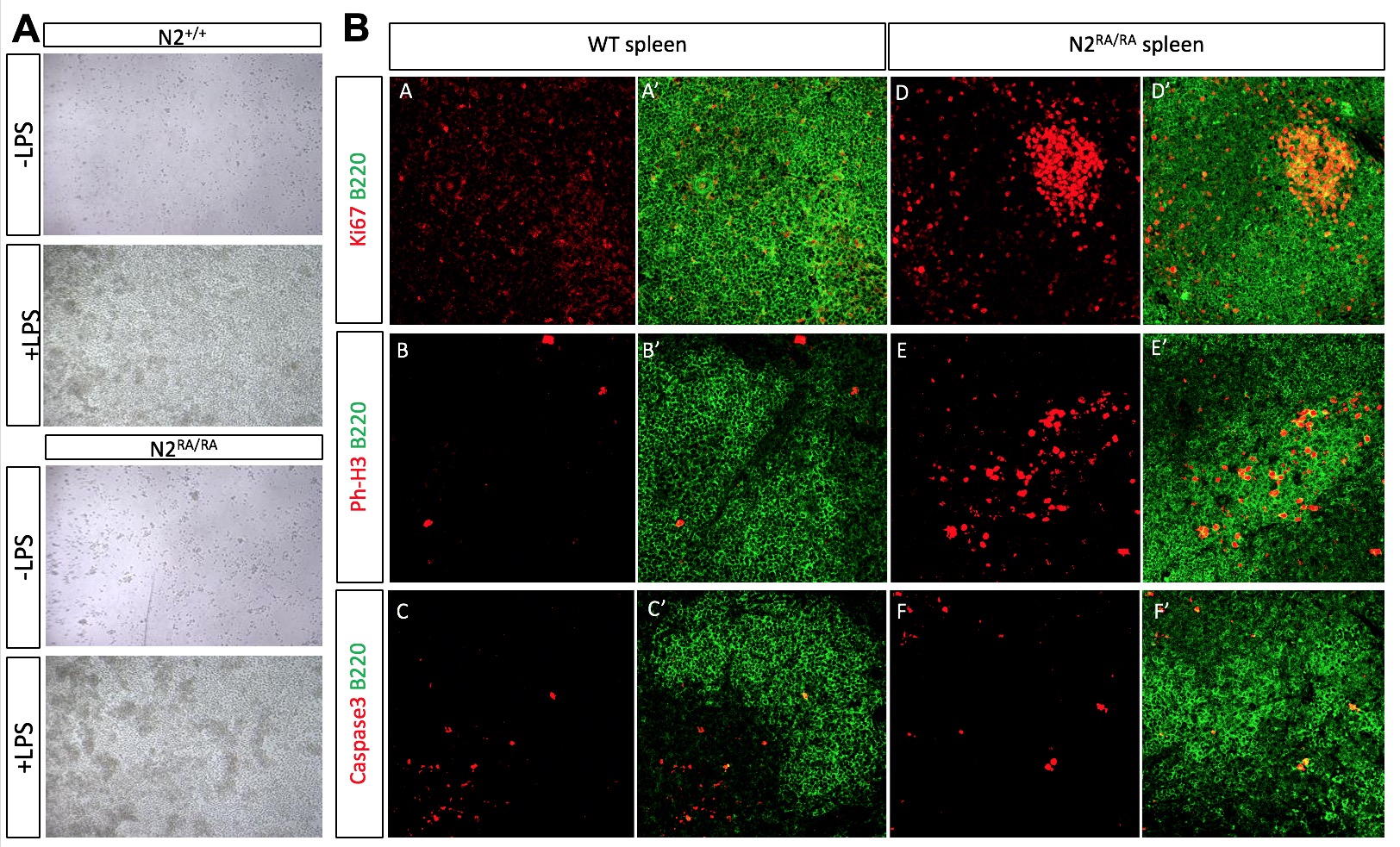

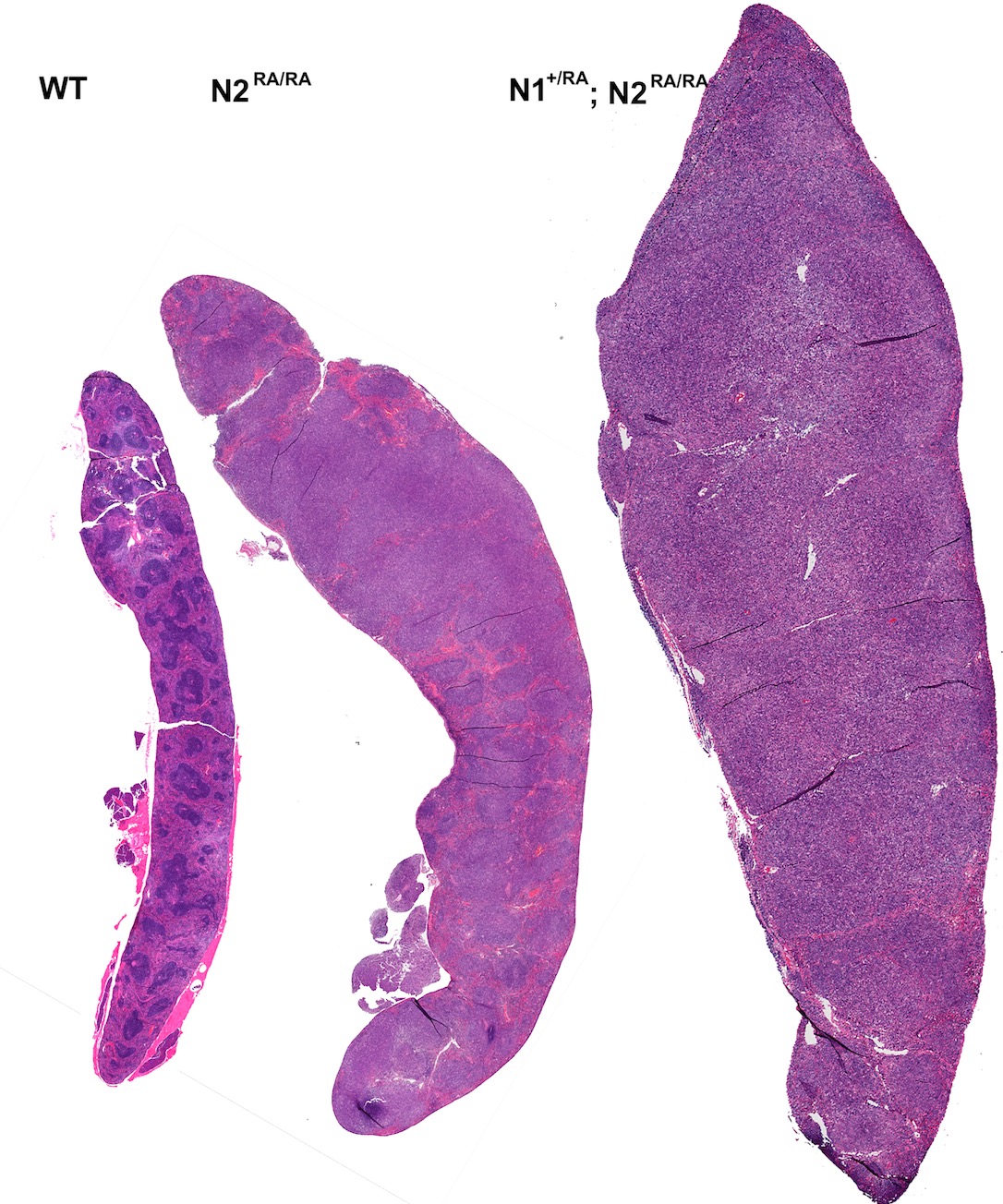
**Supplementary Figure S**
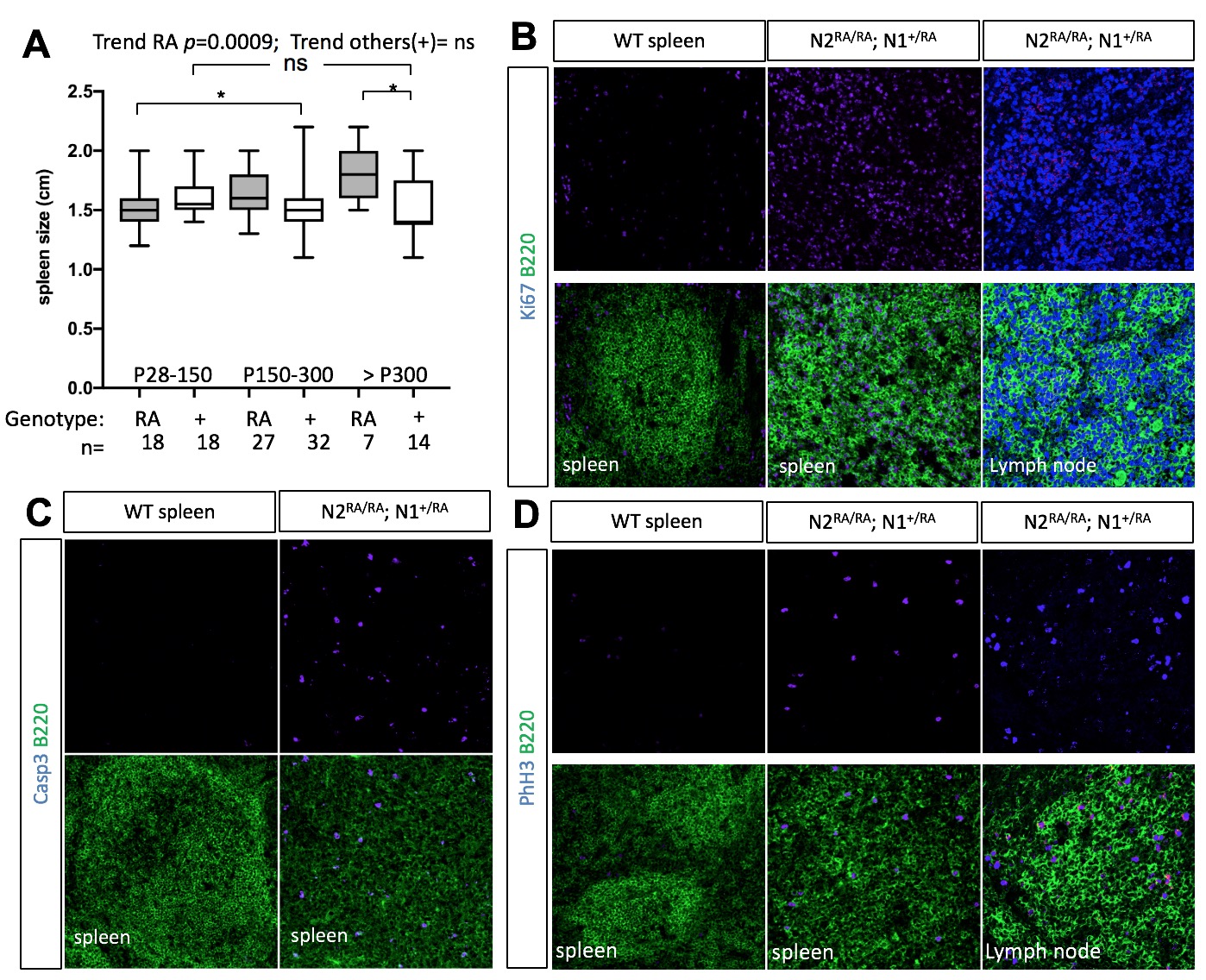
**5:**

**E**

**Supplementary Figure S6:**
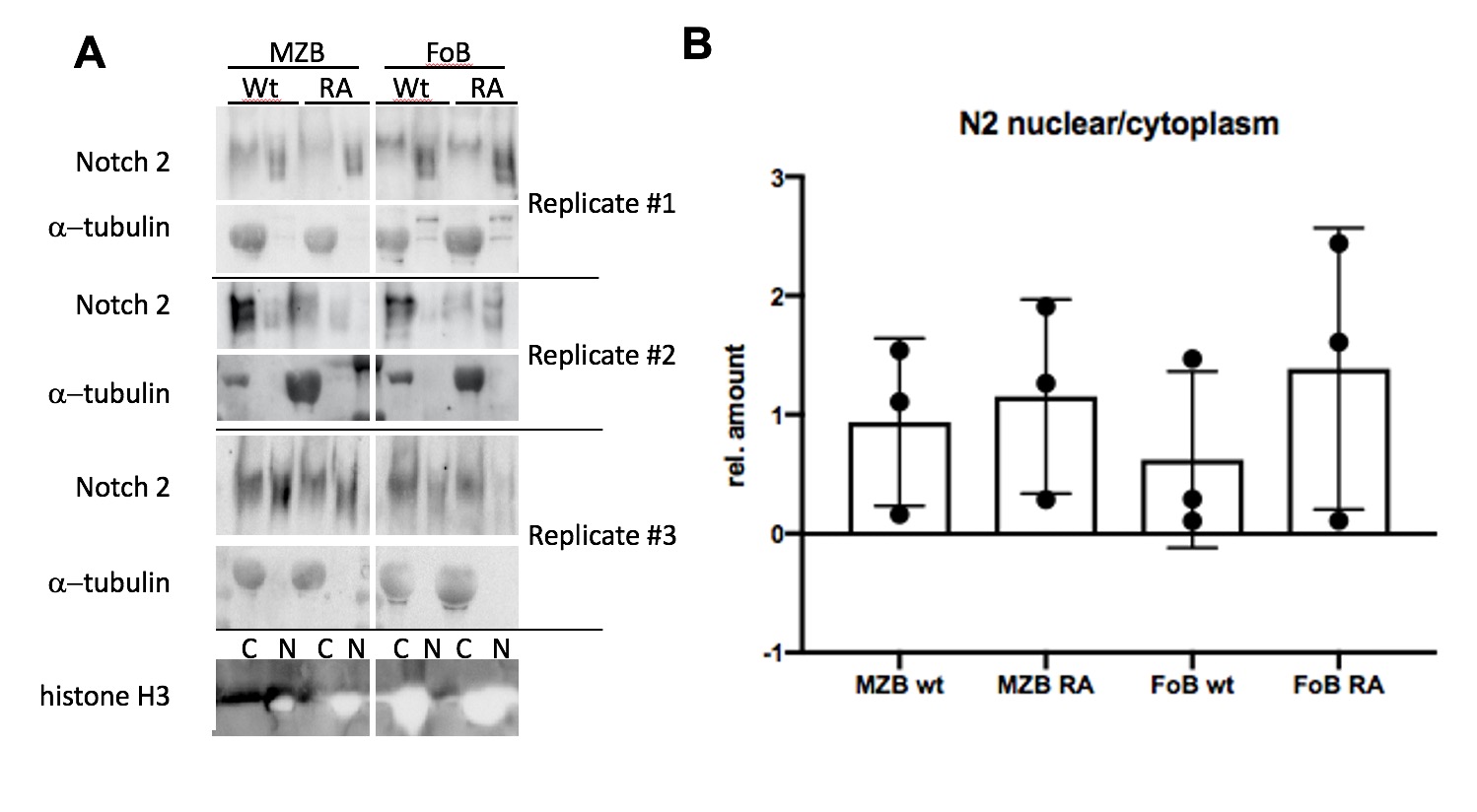

**Supplementary Figure S7:**

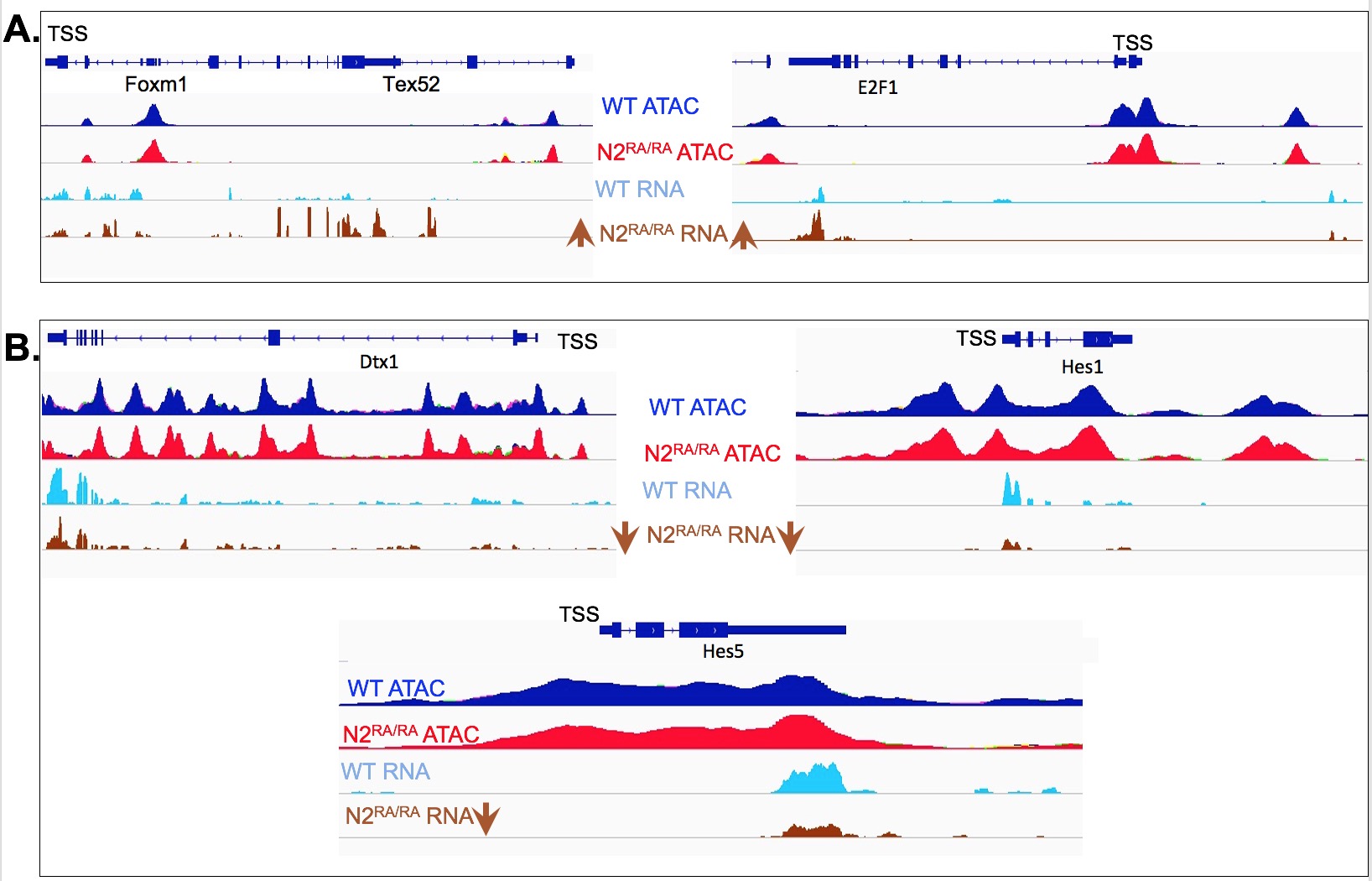

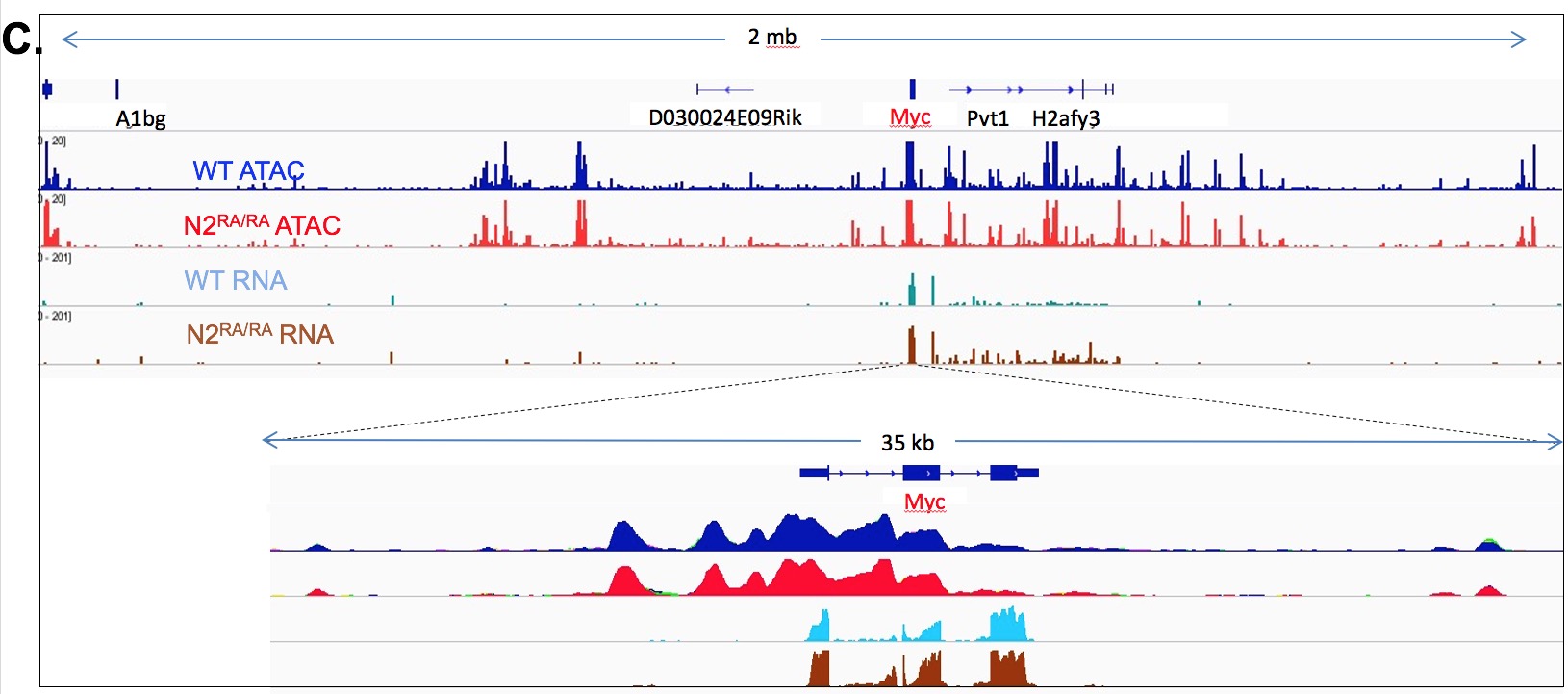

**Supplemental Table S1: Chi-squared analysis of pups (p0 and p1) born in of *N1^RA/RA^;N2^RA/RA^* x *N1^+/–^;N2^+/–^* cross in mite-free housing**

|  | | *expected* | | *observed* | | (O-E)^2^  O |
| --- | --- | --- | --- | --- | --- | --- |
| F1 | *N1^RA/-^; N2^RA/-^* | 25% | 47.5 | 14 | 7.37% | 23.63 |
|  | *N1^RA/-^; N2^+/RA^* | 25% | 47.5 | 47 | 24.74% | 0.01 |
|  | *N1^+/RA^; N2^RA/-^* | 25% | 47.5 | 51 | 26.84% | 0.26 |
|  | *N1^+/RA^; N2^+/RA^* | 25% | 47.5 | 78 | 41.05% | 19.58 |
|  |  | 100% | 190 | 190 | 100% | 43.47 |
| Degree of freedom (# of genotypes -1)=3 | | | | CHISQ.DIST | | ***p*=10^–9^** |

**Supplemental Table S2: Chi-squared analysis of male and female pups born in C57BL/6J *N1^+/RA^* x *N1^+/RA^* cross during fur mite infestation and following generations**

| **Males** | | | | *expected* | | | *observed*  (male/female) | | | (O-E)^2^  O |
| --- | --- | --- | --- | --- | --- | --- | --- | --- | --- | --- |
| F1 | *N1^RA/RA^* | | | 25% | | 14.25 | 12 | 21.1% | | 0.36 |
|  | *N1^RA/+^* | | | 50% | | 28.5 | 33 | 57.9% | | 0.71 |
|  | *N1^+/+^* | | | 25% | | 14.25 | 12 | 21.1% | | 0.36 |
|  | | | | 100% | | 57 | 57 | 100% | | 1.42 |
| Degree of freedom (# of genotypes -1)=2 | | | | | | | CHISQ.DIST | | | ***p*=0.49** |
| **Females** | | | *expected* | | | | *observed*  (male/female) | | | (O-E)^2^  O |
| F1 | | *N1^RA/RA^* | 25% | | 12.5 | | 3 | | 6.0% | 7.22 |
|  | | *N1^RA/+^* | 50% | | 25 | | 30 | | 60.0% | 1.00 |
|  | | *N1^+/+^* | 25% | | 12.5 | | 17 | | 34.0% | 1.62 |
|  | | | 100% | | 50 | | 50 | | 100% | 9.84 |
| Degree of freedom (# of genotypes -1)=2 | | | | | | | CHISQ.DIST | | | ***p*=0.01** |
